# Controlling the human connectome with spatially diffuse input signals

**DOI:** 10.1101/2024.02.27.581006

**Authors:** Richard Betzel, Maria Grazia Puxeddu, Caio Seguin, Vincent Bazinet, Andrea Luppi, Alina Podschun, S. Parker Singleton, Joshua Faskowitz, Vibin Parakkattu, Bratislav Misic, Sebastian Markett, Amy Kuceyeski, Linden Parkes

## Abstract

The human brain is never at “rest”; its activity is constantly fluctuating over time, transitioning from one brain state–a whole-brain pattern of activity–to another. Network control theory offers a framework for understanding the effort – energy – associated with these transitions. One branch of control theory that is especially useful in this context is “optimal control”, in which input signals are used to selectively drive the brain into a target state. Typically, these inputs are introduced independently to the nodes of the network (each input signal is associated with exactly one node). Though convenient, this input strategy ignores the continuity of cerebral cortex – geometrically, each region is connected to its spatial neighbors, allowing control signals, both exogenous and endogenous, to spread from their foci to nearby regions. Additionally, the spatial specificity of brain stimulation techniques is limited, such that the effects of a perturbation are measurable in tissue surrounding the stimulation site. Here, we adapt the network control model so that input signals have a spatial extent that decays exponentially from the input site. We show that this more realistic strategy takes advantage of spatial dependencies in structural connectivity and activity to reduce the energy (effort) associated with brain state transitions. We further leverage these dependencies to explore near-optimal control strategies such that, on a per-transition basis, the number of input signals required for a given control task is reduced, in some cases by two orders of magnitude. This approximation yields network-wide maps of input site density, which we compare to an existing database of functional, metabolic, genetic, and neurochemical maps, finding a close correspondence. Ultimately, not only do we propose a more efficient framework that is also more adherent to well-established brain organizational principles, but we also posit neurobiologically grounded bases for optimal control.

## INTRODUCTION

The human connectome is a network map of the brain’s physical wiring [1, 2]. Nodes correspond to neural elements – brain regions at the macroscale – where computations are carried out locally. The outcomes of those computations are relayed from one node to another along network edges – fasciculated white-matter [3, 4]. Accordingly, the structure of the connectome plays an important role in shaping interregional communication patterns [5] and the correlation structure of brain activity–i.e. functional connectivity (FC) [6].

A node’s state can be defined at any instant based on its magnitude of fMRI BOLD activity (Fig. 1a). Aggregating these values across all brain regions returns an *N* × 1 vector; we refer to this multivariate pattern as a “brain state”, positioning the brain at a specific location in an *N* -dimensional state space at a given time [7]. As the brain’s activity fluctuates over time so does its state, tracing out a trajectory as it transitions from one pattern of activity to another.

**Figure 1.**
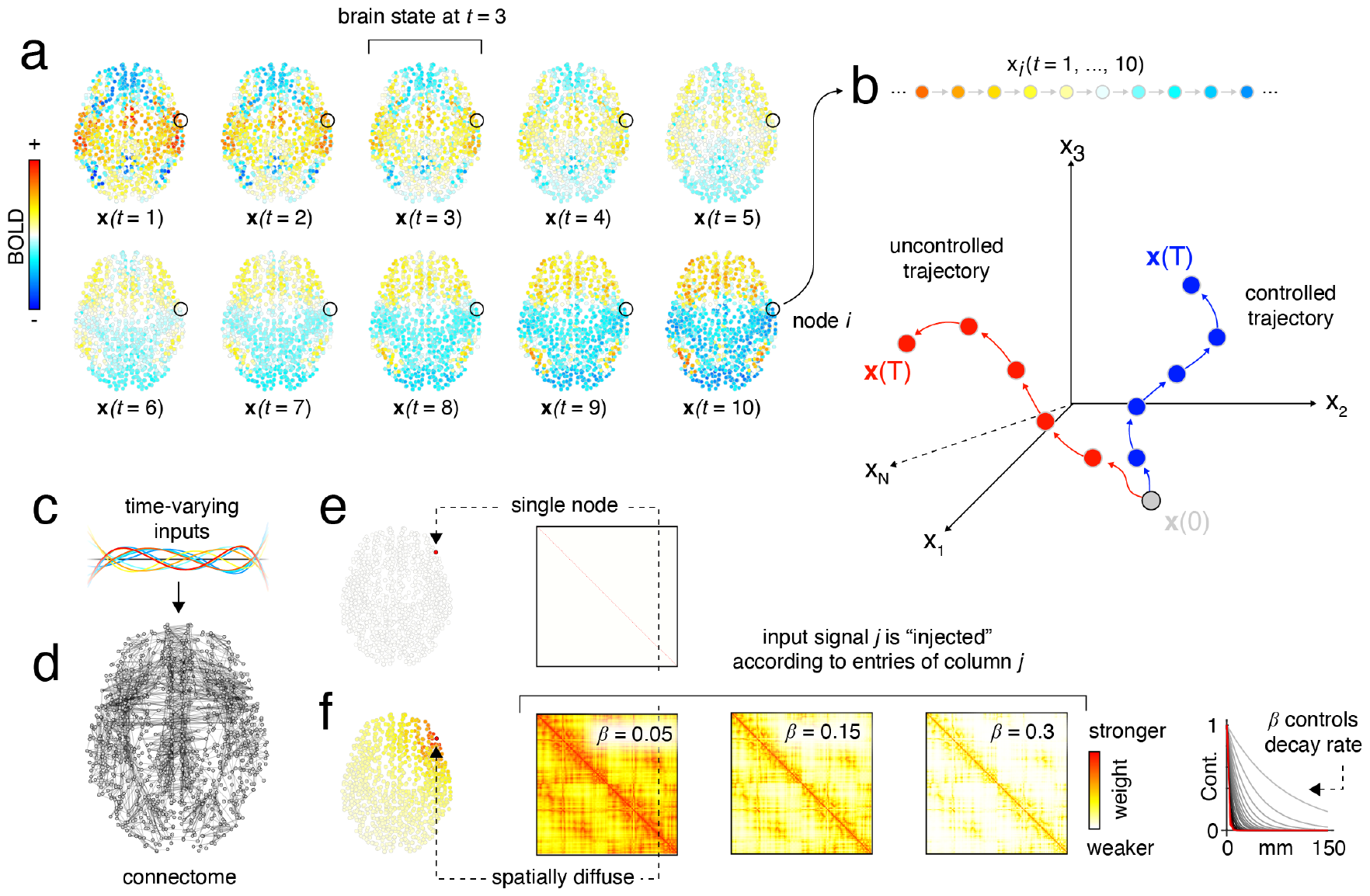
Illustration of optimal control framework. (*a*) Brain activity changes from time point to time point. The activity of region *i* at time *t* is specified as *x*_*i*_(*t*). The set x(*t*) = [*x*_1_(*t*), …, *x*_*N*_ (*t*)] defines the brain’s “state” at time *t*. (*b*) For a given node, *i*, the sequence x_*i*_ = [*x*_*i*_(1), …, *x*_*i*_(*T*)] defines a 1-dimensional trajectory. When we consider *N* possible nodes, we can think of the brain as moving through an *N* -dimensional state space across time, tracing out a high-dimensional trajectory. In the absence of any inputs, the brain will follow an “uncontrolled” trajectory given its intrinsic dynamics. (*c*) However, control signals–time-varying inputs–can be injected into the brain that change local (regional) states and alter the trajectory. (*d*) The signal propagates along white-matter tracts so that the input node relays information about the input to its connected neighbors. The new trajectory is referred to as the “controlled” trajectory. (*e*) Traditionally, each control signal gets injected into a single node. An alternative possibility is that the signal is centered on a particular location in the brain–e.g. a region’s center of mass–but that nearby regions receive some of that signal, with the amount received decaying exponentially with distance as controlled by the by the diffusivity parameter, *β* (see panel *f*).

Even in the absence of input – i.e. experimental stimuli, environmental perturbations, or internally-generated signals – the brain is not static; rather, its state passively evolves along a trajectory owing to the inertia of its own dynamics [8–10]. We refer to this trajectory as “uncontrolled.” It is possible, however, for the brain to deviate from this trajectory were it to receive input.

Recently, there has been considerable interest in investigating whether input signals could be tailored to (deliberately) push the brain along a “controlled” trajectory and into a desired target state (Fig. 1b). This question is presently highly relevant, given the enormous interest in technologies for manipulating ongoing brain activity, including transcranial magnetic [11] and direct current stimulation [12] as well as chemogenetic applications [13] and optogenetic stimulation [14].

One promising framework for understanding the effect of these perturbations is network control theory, a branch of engineering concerned with controlling the behavior of networked dynamical systems [15–20]. The optimal control framework, in particular, has proven especially influential. Briefly, it derives a series of time-varying inputs that push the network into a target state. These inputs are optimal in the sense that they minimize a two-term cost function. The first term refers to the squared amplitude of the input signals integrated over time – the so-called “control energy.” The second term is calculated as the distance from a reference state, typically set equal to either the target state or an *N* × 1 vector of zeros. Here, optimal control balances these two terms, seeking input signals that are low energy while keeping brain state trajectories from straying too far from their reference.

Optimal control has been used to understand the link between brain structure and function [21–24], to study brain network development and executive function [25–27], working memory and schizophrenia [28], epilepsy [29, 30], psychiatric disorders [31, 32], mindfulness training [33], neurostimulation [34, 35], and psychedelic research [36, 37]. Other work has focused on understanding the metabolic cost of control [38, 39].

One of the crucial components of optimal control is the input matrix. This matrix determines the mapping of time-varying input signals (Fig. 1c) to neural elements (nodes) in the brain (Fig. 1d). In most applications, there is a one-to-one mapping of inputs to nodes, such that each input signal is delivered to exactly one node. Though mathematically convenient, this assumption presents some issues. In the context of neurostimulation, it implies that the stimulation technique has perfect precision and specificity. That is, a user is able to target and stimulate a given population of neurons without impacting the states of its proximal neighbors. This is unrealistic and especially so for non-invasive techniques, where not only is the spatial specificity limited, but where errors in target selection may also occur. Additionally, if we were to imagine that the control signal was self-generated by a specific neuronal population, it is not clear that this population will fall neatly within the boundaries of a specific parcel [40]. More likely, the population is distributed across multiple parcels, implying that the input signal is delivered to multiple nodes in the network.

Here, we present a strategy for incorporating spatial dependencies into the optimal control framework. We do so by directly modifying the input matrix to account for nodes’ spatial proximities. Traditionally, to deliver an input to node *i*, we create an *N* -dimensional column vector whose *i*th element is equal to 1. If we wanted to deliver one control signal per brain region, the corresponding control matrix would be the identity matrix (see “single node” example in Fig. 1e). Other studies have modified this matrix, typically by selectively rescaling its elements–e.g. to model the effect of cortical thinning or neurochemical variation [23, 36]. Here, however, we propose a more radical, though eminently simple, modification. Specifically, we deliver every input signal to every brain region–i.e. the control matrix contains all non-zero elements–but scale the weights in the control matrix so that the element *{i, j}* is a monotonically decreasing function of the distance, *D*_*ij*_, of node *j* from the input node, *i* (Fig. 1e,f).

To our knowledge, this type of modification to the input matrix, though anticipated [41], has never been investigated directly. Here, we study the “spatial” input strategy, comparing its performance to that of the more traditional “local” input strategy and using it to develop novel neuroscientific insight. We find that the energy needed to transition between empirically-derived brain states under the spatial input strategy tends to be smaller and gives rise to dissimilar time-varying inputs and regional control energies. We also propose a simple strategy for near-optimal control that takes advantage of correlations among input signals–a measure of compressiblity–to further reduce the amount of energy on a per-transition basis by orders of magnitude. Finally, we show that the topography of the optimal input patterns is correlated with well-established transcriptomic, neurochemical, metabolic, and functional brain maps. Though simple, the extension proposed here pushes the realism of the optimal control framework and opens up avenues for future studies to link control with empirical observations.

## RESULTS

Here, we compare “local” and “spatial” strategies for delivering control signals. The local strategy injects each input into one brain region and one region only, whereas the spatial strategy allows for those input signals to diffusely impact the states of nearby regions. We apply these input strategies to a multi-modal human imaging dataset that includes both diffusion spectrum and resting-state imaging data (*N*_*s*_ = 70 adults age 28.8*±*9.1 years; 43 males) [42]. We replicate our main findings using Human Connectome Project data (*N*_*s*_ = 95) [43].

### Defining brain states

The optimal control framework considers transitions between discrete brain states – i.e. whole-brain patterns of activity. Before we can determine the optimal input signals, we must first define brain states. Unlike previous studies, which have defined states meta-analytically [23] or based on canonical large-scale systems [21, 22], we do so empirically using the following algorithm. First, we detect peaks in the amplitude of whole-brain activity (root mean square across *N* = 1000 regions/nodes [44]). We define peaks as frames whose RMS was greater than that of both the preceding and following frames. We then aggregate the corresponding patterns (*N*_*peak*_ = 3443 patterns in total) across participants and calculate Lin’s concordance between all pairs of patterns. We then cluster the resulting *N*_*peak*_×*N*_*peak*_ matrix using a variant of modularity maximization [45, 46]. The algorithm returned 139 clusters. Most were small and/or contained peaks from only a few participants. We defined brain states as the centroids of clusters in which peak activations from at least 50% of participants were represented (see **Materials and Methods** for more details). Note that this approach for empirically defining brain states is similar to that of [25] and [36], wherein fMRI BOLD time series were clustered into recurring states. It also resembles the well-established method for extracting co-activation patterns (CAPs) [47–49] from fMRI BOLD data (though note here that our approach will include low-amplitude peaks, whereas CAPs retains more extreme peaksx).

This procedure resulted in *N*_*state*_ = 11 brain states (Fig. 2a) that appeared in anywhere from 99% to 51% of participants. Broadly, these brain states recapitulated well-known activation patterns, delineating previously described large-scale brain systems [50] (Fig. 2b). For instance, State 1, which appeared in 99% of all participants, corresponded to activation of regions in the salience/ventral attention and somatomotor networks and deactivation of regions in the default mode and control networks. State 2 appeared in 99% of participants, and is almost perfectly anti-correlated with State 1, corresponding to default mode activation and salience/VAN and somatomotor deactivation. State 3 corresponds to activation of default mode and control regions; states 4 and 5 are mirrors of one another and correspond to activation/deactivation of visual regions, respectively; state 6 corresponds to activation of dorsal attention and control regions and deactivation of the default mode; state 7 corresponds to activation of the somatomotor network and deactivation of the control network; state 8 corresponds to deactivation of the visual network; state 9 corresponds to activation of the control network; state 10 corresponds to activation of default mode regions and deactivation of dorsal attention regions; state 11 corresponds to deactivation of the control network (see Fig. 2a,b,e for topographic maps of each state, their association with brain systems, and their correlation with one another, respectively).

**Figure 2.**
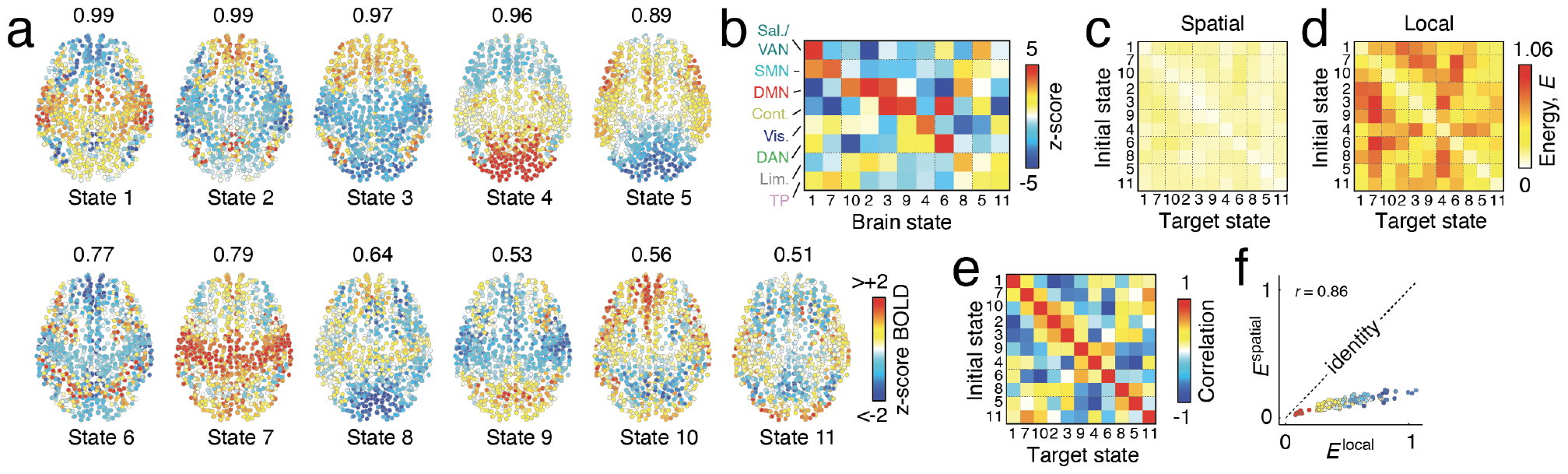
Brain state transitions. (*a*) Spatial topography of eleven empirically-derived brain states. Text above each brain represents the fraction of participants in which a given brain state was observed. Note that we disallowed states that appeared in fewer than 50% of participants. (*b*) z-scored overlap between brain states and canonical resting-state networks. For each brain state and brain system, we calculate the mean activity within that system. We then compare the mean activity against a space-preserving null model and express the mean similarity as a standard score (z-score). Large positive values indicate that the mean activity of nodes within a given system is greater than expected by chance. Large negative values indicate that the mean deactivation of nodes is greater (more negative) than expected by chance. (*c*) Global energy associated with the transition between all pairs of brain states under the “spatial” model in which the control input centered on region *i* is also delivered to region *i*’s neighbors with an amplitude inversely proportional to the Euclidean distance from region *i* (negative exponential). (*d*) Global energy associated with the transition between all pairs of brain states under the “local” model in which a control input is delivered to region *i* alone. (*e*) Spatial similarity of brain state activations with respect to one another. (*f*) Scatterplot comparing the energy associated with every possible transition under both input models. The color points corresponds to their pairwise similarity from panel *e*.

Next, we calculated the energy associated with transitioning between all pairs of states (121 transitions in total) as the mean squared amplitude of input signals over time. We performed these control tasks separately for the “local” and “spatial” input strategies, arbitrarily fixing the diffusivity parameter to *β* = 0.15 (see Fig. 2c,d for whole-brain energies associated with all transitions). Note that in the following subsections we characterize performance as we vary the value of *β*.

The diffusivity parameter controls the extent to which an input signal centered over node *i* impacts nodes in the immediate spatial neighborhood. One can think of the spatial diffusivity in two contexts. If we imagine that the control signals are being delivered exogenously, e.g. *via* stimulation, then the diffusivity of inputs might reflect the spatial resolution/specificity of the TMS signal (on the order of 1 cm [51–53]) or volume conduction [54]. On the other hand, if we imagine that control signals are delivered endogenously, then the spatial diffusivity could reflect either propagation of electrical signals *via* superficial fibers [55] that may be poorly reconstructed from dMRI and tractography data [56]. Smaller and larger value of *β* correspond to larger and smaller neighborhoods, respectively. We then compared global control energies between local and spatial strategies for each of the 121 possible transitions. Although the energies were correlated with one another (*r* = 0.86, *p <* 10^−15^; Fig. 2f) and related to the spatial similarity of brain states (*r* = −0.82, *p <* 10^−15^; Fig. 2e), we found that the energy associated with the spatial strategy was significantly less than that of the local strategy (paired sample *t*-test; *t*(120) = −22.6, *p <* 10^−15^). Note that we replicated this finding using Human Connectome Project [43] data (*t*(143) = −22.7, *p <* 10^−15^; Fig. S1). We also show that this result replicates using alternative definitions of brain state (Fig. S2) and with different parcellations (Fig. S3) and as we vary the free parameter, *ρ*, in the optimal control algorithm (Fig. S4).

Together, these results hint that the spatial input strategy may be less effortful than the local strategy. It is unclear, however, whether there are other distinctions between the two strategies and how much these observations depend on the precise parameterization (the value for *β*) of the spatial strategy.

### Spatially diffuse control inputs reduce control energy and correspond to distinct state trajectories and input signals

In the previous section, we demonstrated that on a per-transition basis, when the spatial diffusivity parameter was set to *β* = 0.15 the spatial input strategy required less effort (reduction of the mean squared control inputs averaged across all brain regions) compared to the local strategy. Note that the value of *β* = 0.15 corresponds to, approximately, the widest gap between transition energy for the spatial and local input strategies. Here, we investigate how these properties vary as a function of the spatial diffusivity parameter, *β*, whose value we limited to the interval [0.01, 0.5]. The lower bound of this range was selected because smaller values led to instabilities in the estimation of optimal inputs; the upper bound was selected as values beyond *β* ≈ 0.5 yielded results that were qualitatively indistinguishable from smaller values of *β*. We focus on addressing four specific questions.

Here, optimal control seeks to minimize a cost function that balances distance from the target state with the squared amplitude of control inputs, integrated over time (this latter term is referred to as the control energy). Our first and second questions relate to the relative contributions of these two terms. First, does the spatial strategy change the character of the dynamic trajectory? To address this question, we calculated the mean distance from the target state across the entire controlled trajectory under both the spatial and local strategies (Fig. 3a). Then, for a given transition and *β* value, we calculated the difference in distance between the spatial and local models. Interestingly, we found that for small values of *β*, which correspond to greater spatial diffusivity, the difference in distance was large, indicating that the spatial model produced trajectories towards the target state that were less direct (Fig. 3d). As *β* increased and the inputs approximated that of the local strategy, the gap narrowed. See Fig. S5 for example trajectories.

**Figure 3.**
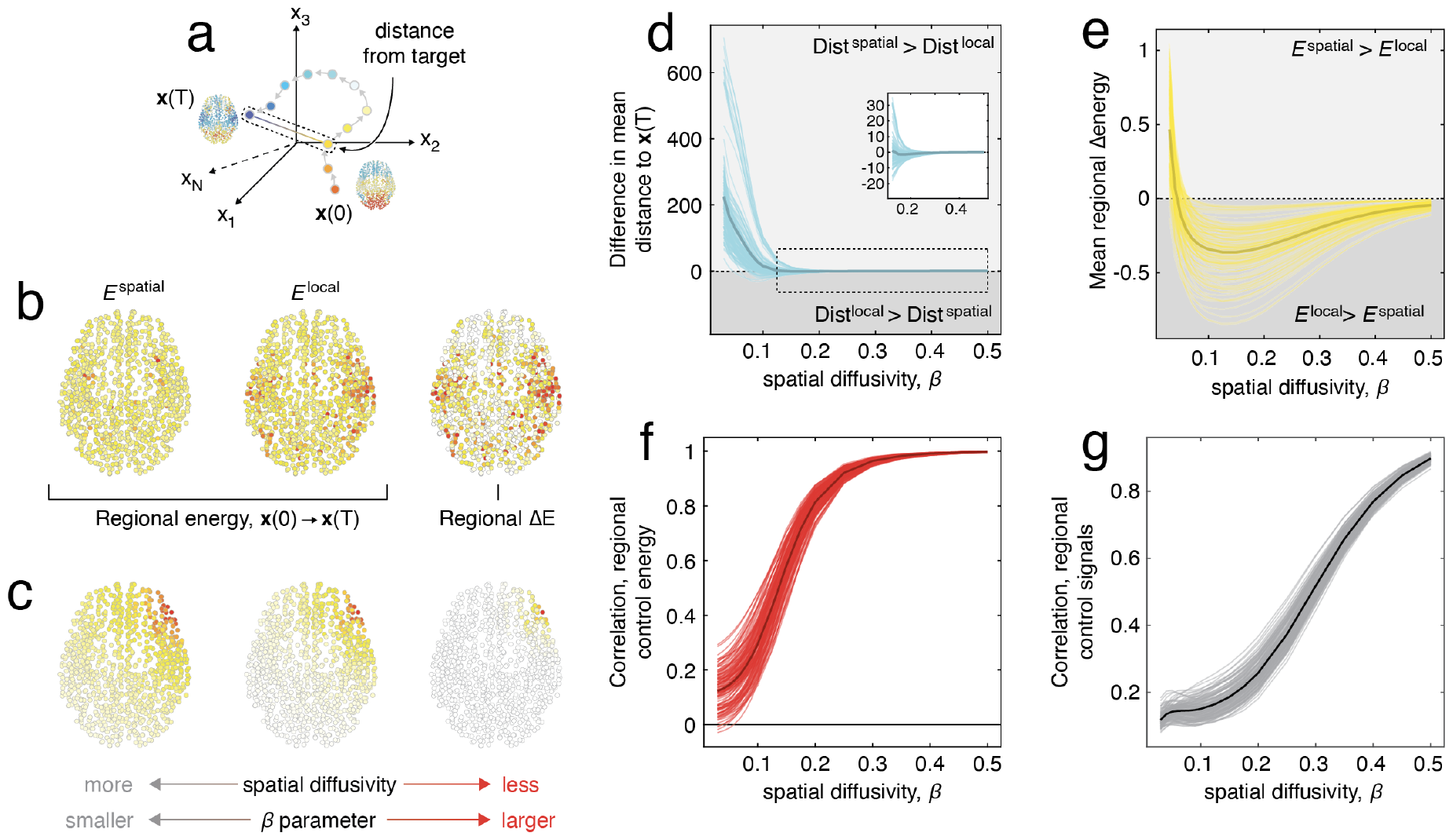
Comparison of local and spatially diffuse control strategies. (*a*) We consider transitions from an initial state x_0_ to a target state x_*T*_ . For each such transition, we measure the mean distance from the target state to each point along the controlled trajectory. (*b*) We calculate regional control energies for both the “spatial” and “local” input strategies. Using those values, we derive two additional metrics: the average difference in regional energy and the correlation of the *N* × 1 vector of regional energies. We also calculate, for every brain region, the correlation between the input signals centered on that region under “spatial” and “local” input strategies. (*c*) We repeat these calculations as we vary the spatial diffusivity parameter, *β*, from a highly diffuse regime (small values of *β*) to a spatially focused regime (larger values of *β*).(*d*) Mean distance from target state as a function of *β*. Each trajectory represents one of 121 possible transitions. The inset highlights 0.125 *< β <* 0.5. (*e*) Difference in regional control energy averaged across all *N* = 1000 cortical regions of interest at each *β* value. (*f*) Spatial similarity (correlation) of regional control profiles. (*g*) Temporal similarity (correlation) of regional control inputs.

Second, we asked the complementary question: does the effort needed to perform control tasks vary with *β*? To address this question, we calculated the mean squared difference in regional energy between the local and spatial control strategies, yielding a single scalar value (Fig. 3b). We found that, with the exception of very small values of *β*, the spatially diffuse inputs generally required less energy (Fig. 3e). Interestingly, the gap between spatial and local control energy increases as the similarity between input and target states grows (this is evident in Fig. 2f). In the supplementary material, we also confirmed that this difference is not obviously related to differences in the total weight of the input matrix (Fig. S6b and Fig. S7).

Third, we asked whether the *regional* control energies – the mean squared amplitude of the control signal delivered to each region – varied with *β*. That is, how does the multivariate pattern of input energy across the brain change as we vary the diffusivity of input signals? To do this, we estimated the *N* × 1 vector of regional control energies for each transition at every *β* value, and calculated its similarity with respect to the regional control energy vector obtained under the local input strategy (Fig. 3f). Again, we found that, for small values of *β*, the control patterns were dissimilar, but with similarity monotonically increasing as a function of *β*.

Finally, we asked whether the optimal input signals associated with the spatial and local input strategies were correlated across time (see Fig. S8). That is, are the time-varying inputs – rather than their mean amplitude – similar between spatial and local strategies and does that level of similarity vary with *β*? Analogous to control energy, we found that temporal similarity increased near monotonically with *β*, though at a slower rate and never fully saturating over the range of *β* explored here (Fig. 3g).

Collectively, these results suggest that the spatially diffuse input strategy can yield distinct brain state trajectories, corresponds to reductions in the energy/effort needed to complete control tasks, and does so with distinct sets of control inputs. It also suggests that in the limit as *β* → ∞ the spatial strategy becomes indistinguishable from the local strategy, as expected.

### Differences in local control energy

In the previous section we focused on *global* differences in optimal control as a function of the diffusivity parameter, *β*. Here, we fix the parameter to a value *β* = 0.15 and examine differences at the *local* (regional or nodal) level, focusing on energies and three quantities of interest. First, we calculate the regional control energy under both the local and spatial strategies. We also calculate the *effective* control input to each brain region under the spatial strategy as sum of its neighbors’ inputs weighted by the elements of the input matrix, i.e. **u**^*eff*^ = **B**^*eff*^ **u**^⊺^, where **B**^*eff*^ = *exp*(−*β ·* **D**). From the effective input, we then calculate the effective energy at region *i* as 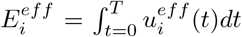. The effective input reflects the incidental amount of energy delivered to each node given its geometric relationship to other nodes in the network.

In general, we found significant correlations between all three measures (*r*_*local,spatial*_ = 0.58 *±* 0.06; *r*_*local,eff*_ = 0.59*±*0.10; *r*_*spatial,eff*_ = 0.75*±*0.02 Fig. 4a-c); this effect held over most ranges of *β* (Fig. S6a). However, and as in the previous subsection, the regional energies were lowest for the spatially diffuse inputs (one-way ANOVA, *F* (2) = 27692.2; *p <* 10^−15^; post-hoc paired-sample *t*-tests between spatial and effective, effective and local, and spatial and local had minimum *t*(120999) = 41.2; all *p <* 10^−15^).

**Figure 4.**
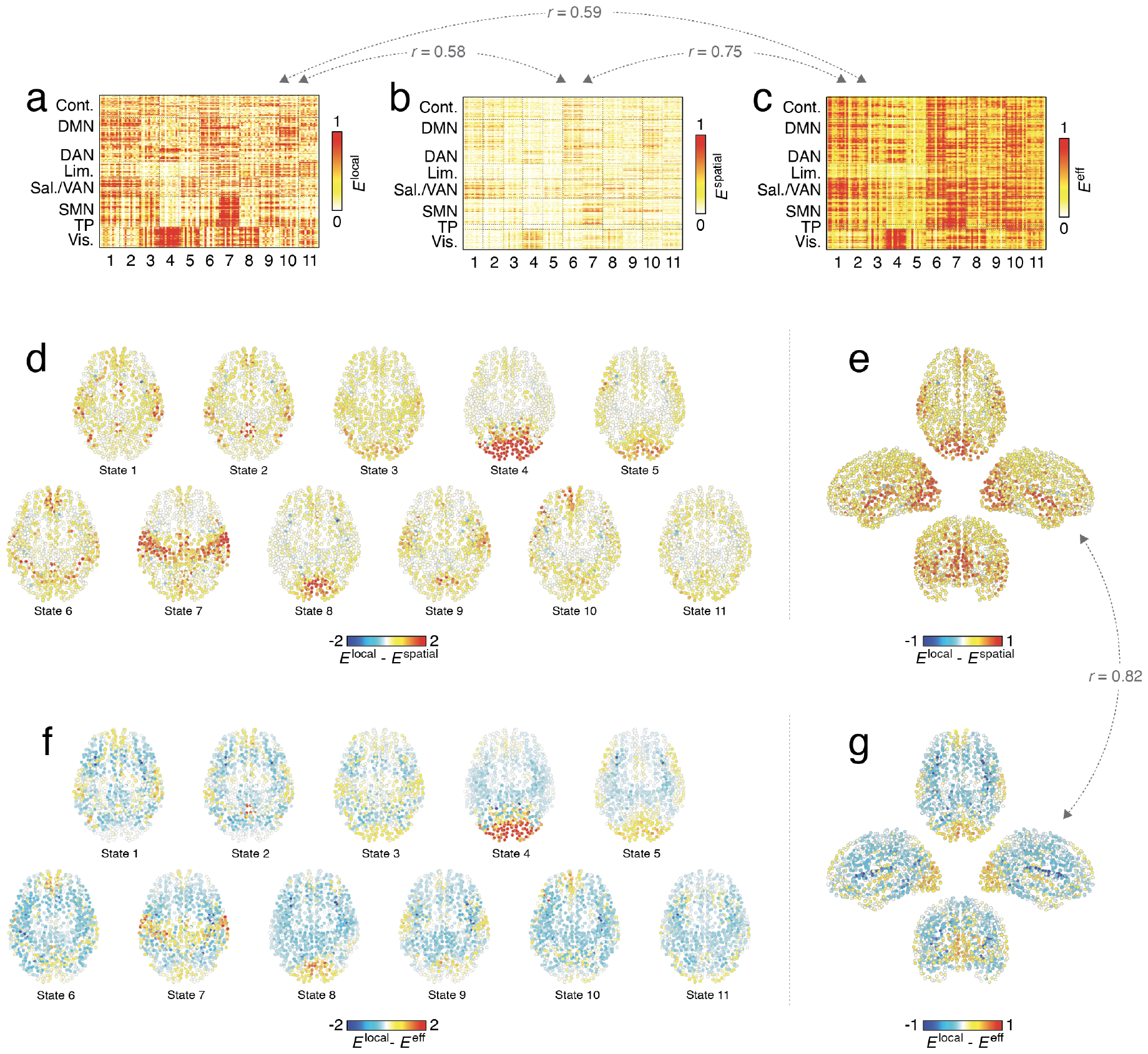
Comparing regional control energies. Panels *a, b*, and *c* show the “local”, “spatial”, and “effective” input energies at each brain region and for each of the 121 control tasks. Note that the columns in each matrix are ordered based on target state. The gray text above the heatmaps indicates the mean correlation between columns of these matrices. (*d*) Differences in control energy (local minus spatial). Warm colors indicate regions where the control energy under the local strategy exceeded that of the spatial strategy. Note that for brain plots, the regional differences in energy are averaged over all transitions in which a given state was the target. For example, the “State 1” sub-panel corresponds to the mean difference in energy for transitions, 2→1, 3→1, 4→1, and so on. Panel *e* shows the mean difference across all target states. Panels *f* and *g* are analogous to *d* and *e*, but compare local input strategy with the effective input strategy.

These results suggest that the different input strategies correspond to similar patterns of regional effort but of dissimilar energy. How are these energy differences distributed spatially? Is it the case that all nodes uniformly increase/decrease their energy from condition to condition or are the differences more focal? To test this, we calculated the difference in regional control energies for local *versus* spatial (Fig. 4d,e). We found that the differences were highly dependent on the the topography of the target state. Regions whose target activity was very positive or very negative required significantly more energy to activate/deactivate under the local control strategy than the spatial strategy, whereas regions with relatively low-amplitude activity were largely the same when comparing across input strategies (Fig. S9). Interestingly, most regions exhibited reductions in energy under the spatial input strategy. However, a small number of regions actually exhibited increases in energy–i.e. *E*^*local*^ *< E*^*spatial*^ (Fig. S10). These regions tended to overlap across state transitions, were concentrated in the salience/ventral attention network, and correlated with specific neurochemical profiles, including dopamine and acetylcholine transporters (DAT and VAChT, respectively).

We also wanted to assess whether the effective energy associated with the spatially-diffuse inputs differed from that of the local input strategy. For example, it could be the case that the effective energy is similar to that of the local strategy–i.e. the spatially-diffuse inputs take advantage of the brain’s geometry to effectively approximate the optimal inputs under the local policy. In general, and in line with the previous findings, we discovered that highly active/inactive regions required greater energy under the local model compared to the effective input energy (Fig. 4f,g). There was, however, one notable distinction. Specifically, regions with weak activity (close to zero) exhibited slightly reduced energy under the local policy compared to the spatial (Fig. S9).

Altogether, these results indicate that the patterns of regional control energies of the proposed (spatial) model are consistent with those of the conventional (local) one. However, under the spatial model the energy required for the state transition is lower, above all in brain regions with high-amplitude target activity.

### The compressibility and low-dimensional structure of input signals

To this point, we have considered “full control” transitions, whereby input signals are delivered to every node in the network (or in the case of the spatial strategy, the input signals are positioned over every node but allowed to influence neighboring nodes as well). However, visual inspection suggests that many of these input signals are highly correlated and nearly identical, suggesting that the effective dimensionality of the control input is far smaller than the *N* input signals currently being delivered [57]. This prompts the question: Could we ever recover the effective number of input signals? If we could construct archetypical input signals based on this low-dimensional representation, how well would these signals approximate optimal control? That is, would small deviations in the input signal result in large errors so that the system ends up far from its desired target state? In this section, we explore these questions.

For a given control task in which input signals are delivered through *N*_*inputs*_ input sites, we obtain *N*_*inputs*_ time-varying control signals. In practice, these signals can be grouped into clusters (Fig. 5a,b). Each cluster is associated with its corresponding centroid (the mean control signal across all nodes assigned to that cluster), and therefore a representative control signal. Rather than deliver control signals to each node separately, we can imagine delivering this common signal to the nodes assigned to the corresponding cluster, modifying our input matrix, **B**, accordingly (Fig. 5c). This has the effect of reducing the total number of unique inputs from the number of nodes, *N*, to the number of clusters, *k*, while still delivering control signals to every node. This procedure will also generate a map dictating to which nodes the cluster centroid should be delivered (Fig. 5d; see also Fig. S11 for “dominance” maps derived from the input maps). However, this reduction in dimensionality is accompanied by increased error; because the representative control signals will always be imperfect approximations of the true optimal inputs, they will drive the brain into a configuration that is also only an approximation of the desired target state (see Fig. S12 for an example showing how dimensionality reduction interacts with error in target state).

**Figure 5.**
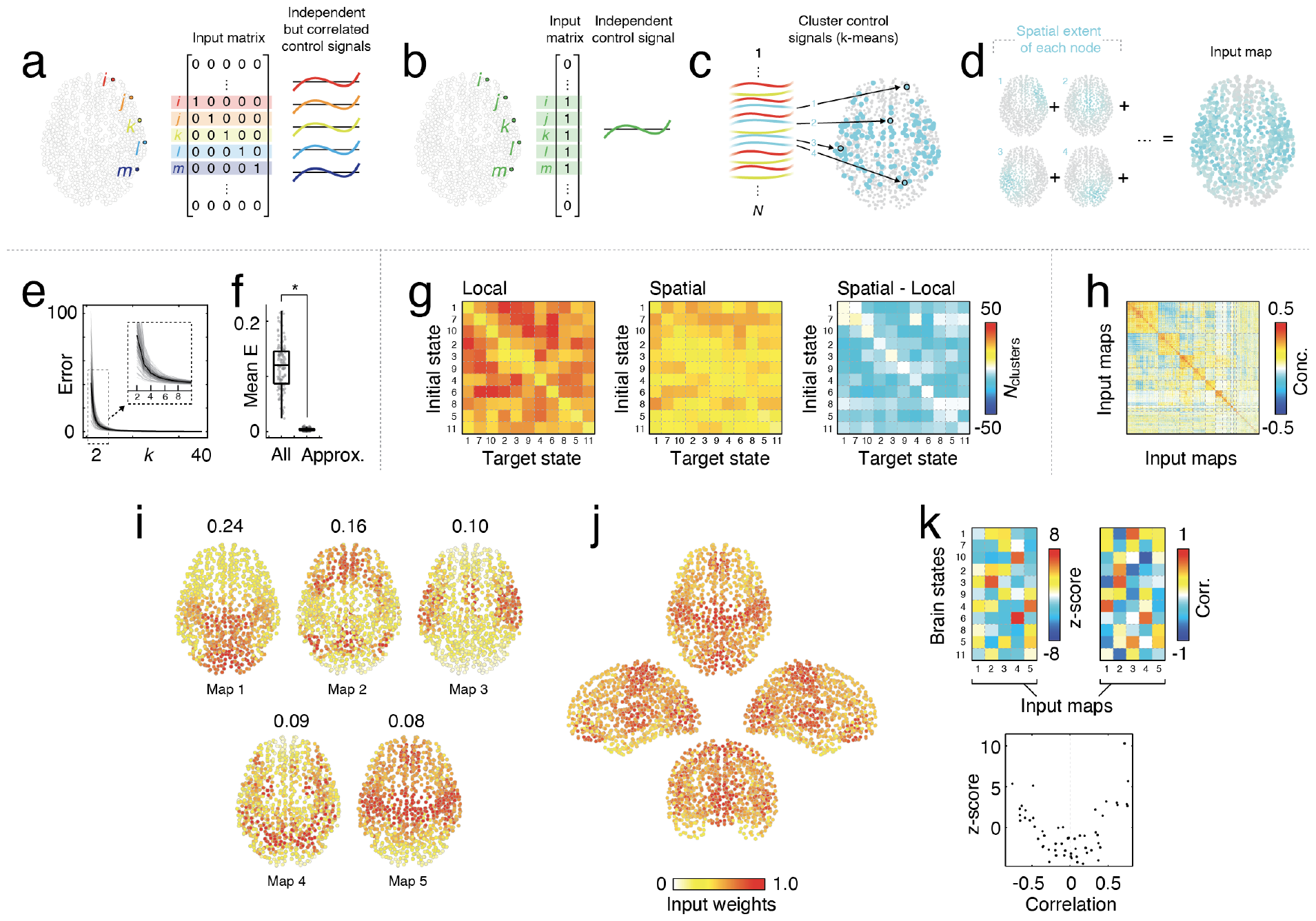
Approximating optimal control with few input signals. (*a*) When many regions deliver input to the brain, their input signals tend to be correlated. (*b*) We can achieve an identical effect by modifying the input matrix so that a single representative input signal gets delivered to multiple regions in the brain. (*c*) Here, we use a *k*-means clustering algorithm to partition input signals into clusters and identify every node associated with a given cluster (as an example, we show the *cyan* cluster here). (*d*) We then obtain a composite map of that cluster by summing the corresponding columns of the input matrix. Then, the cluster centroid (its representative input signal) can be delivered through the composite input map. We vary the number of clusters from *k* = 2 to *k* = 40. We measure how close we get to the target state using the cluster centroids as input signals. (*e*) Error in target state as a function of the number of clusters, *k*. (*f*) Mean energy associated with transitions. (*g*) For a given transition, we can ask on average how many clusters we need before the error falls below a specific threshold. The *left* and *center* heatmaps indicate the number of clusters needed to achieve an error of *ε* = 1*/*1000. The *right* heatmap shows their difference; cool colors are transitions where the spatial model requires fewer clusters. (*h*) We can combine input maps across tasks and cluster them. Here, we show the pairwise concordance matrix ordered by clusters. (*i*) Top five centroids by cluster size. The values above each brain represent the fraction of all input maps assigned to that given cluster. (*j*) The maximum input weight for each region across the top five maps. (*k*) We calculated the similarity of the input maps to each of the eleven brain states involved in the transitions. We found that the input maps were “enriched” for particular brain states (greater similarity than expected) but with no clear one-to-one mapping.

In general, we found that even with relatively few clusters – i.e. small *k* – we could obtain good approximations of the optimal strategy, as measured by the difference in state of the system at time *T* from the target state (Fig. 5e). Notably, this strategy reduced the total control energy by a factor of 38.1 *±* 17.1; that is, the independent control strategy was 3800% more effortful than that of our approximation (Fig. 5f).

For the sake of comparison, we also asked how many clusters–i.e. dimensions–are needed to achieve the same error (deviation from target state) using the local *versus* spatial input strategies. Specifically, we identified the fewest number of clusters, *k*, at which the mean node-level error was ≤ 1*/*1000. In general, we found that the local strategy tended to require larger values of *k* compared to that of the spatial model (5.1 *±* 3.9 more clusters on a per-transition basis; paired-sample *t*-test, *t*(120) = 16.1, *p <* 10^−15^; Fig. 5g).

As noted earlier, this approximate version of optimal control yields brain-wide input maps corresponding to how much of each cluster centroid–a representative control signal–should be delivered to any given node. We aggregated these maps across all 121 control tasks, yielding 1711 maps in total (note that the number of maps varied between tasks, as different numbers of inputs–i.e. clusters–were required to achieve comparable error in their respective target states). We then clustered these maps using the same algorithm previously used to estimate brain states (Fig. 5h). In Fig. 5i we show maps corresponding to the centroids of the five largest clusters, collectively accounting for 67.1% of the total number of maps. We also show the mean input map (Fig. 5j). The input maps occur disproportionately across brain state transitions; some maps show clear preferences for transitions involving particular brain states. For instance, maps 2, 4, and 5 tend to appear in transitions involving brain state 3, brains states 10 and 6, and brain state 4 (Fig. 5k, *top left*). Though others show a less clear preference; the distributions of maps 1 and 3 across brain states is more homogeneous. In general, the tendency for maps to be associated with transitions into a brain state is related to the spatial similarity of the map with that state (Fig. 5k, *bottom*).

These results suggest that we can obtain low-energy approximations of optimal control by driving multiple regions with the same control input. Although the same approximation algorithm can be applied to the local and spatial input strategies, we find that the spatial strategy requires fewer inputs and lower energies to achieve an equivalent approximation of optimal control.

### Input maps align with brain annotations

In the previous section we demonstrated that it was possible to approximately control the brain using a small set of inputs that were coupled, selectively, to a large number of regions. This procedure generates a series of brain maps that represent the coupling of inputs to regions. We showed that although these maps have a link to the topography of initial/target states, the mapping is inexact, prompting the question: What other brain annotations are the input maps linked to? In this section, we address this question by leveraging a large and recently published set of brain annotations [58].

We calculated the correlation between six input maps – the top five maps in terms of cluster size and the maximum input weight across the five maps – with 57 annotations representing neurochemical receptor densities, metabolic measures, brain structural and functional metrics, and spatial transcriptomics (Fig. 6a) [59]. We compared the observed correlation magnitudes with those obtained under a space-preserving null model– the so-called “spin test”–and identified those correlations whose magnitude was significantly greater than chance (accepted false discovery rate fixed at *q* = 0.05; adjusted critical value of *p*_*adjusted*_ = 8.57 × 10^−4^; Fig. 6b).

**Figure 6.**
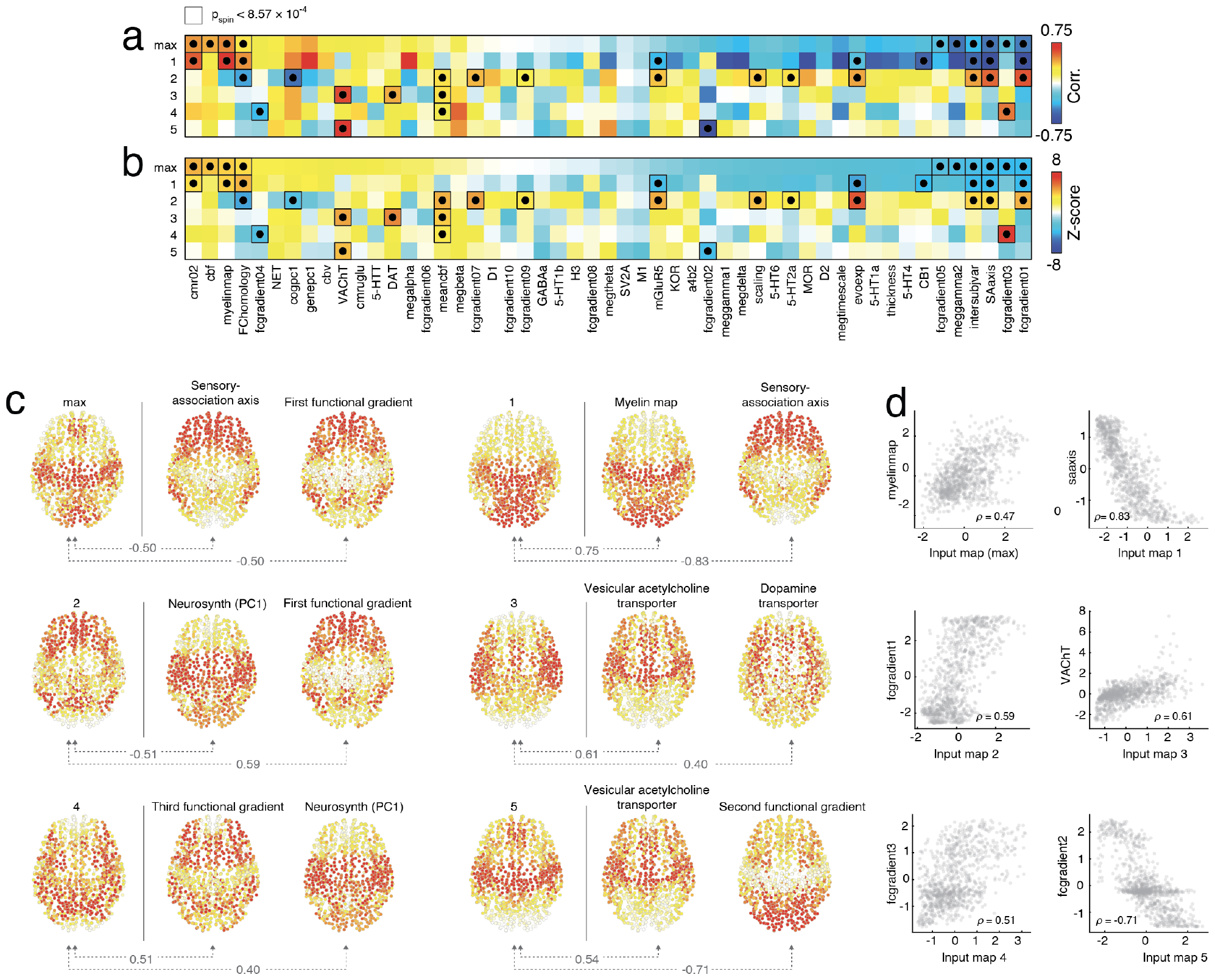
Comparing input maps to brain annotations. (*a*) We calculated the correlation between six input maps with 57 additional maps corresponding to metabolism, neurochemical densities, brain function, brain structure, and spatial transcriptomics. (*b*) We then compared each observed correlation against a null distribution generated under a “spin test”, yielding z-scores for each correlation coefficient, which we used to identify statistically significant correlations. (*c*) Each input map alongside the annotated maps with the strongest correspondence and that passed statistical testing. (*d*) Scatterplots depicting the relationship between select sets of input maps and annotated maps.

We identified multiple statistically significant associations of input maps to annotations. In Fig. 6c,d we show a select set of the strongest associations for each input map alongside scatterplots. Input map 1, which was the largest in terms of cluster size, was strongly correlated with the sensory-association axis [60, 61] (SAaxis; *ρ* = −0.83) and brainwide myelination (*ρ* = 0.75). Other maps exhibited strong associations with cognitive and functional measures. For instance, input map 2 exhibited a close correspondence with the first principal component of the Neurosynth cognitive atlas [62, 63] (cogpc1; *ρ* = 0.55) and the first functional gradient [64] (fcgradient1; *ρ* = 0.59). Other input maps were significantly correlated with neurochemical receptors, including maps 3 and 5, which were correlated with vesicular acetylcholine transport (VAChT; *ρ* = 0.61, *ρ* = 0.54, respectively), while map 3 was correlated with dopamine transport (DAT; *ρ* = 0.40).

Collectively, these observations complement other recent studies seeking a neurobiological basis for the effort measured using optimal control [23, 38].

## DISCUSSION

Optimal control is an engineering framework that has recently been applied in neuroscience, where it has become useful to studying brain state transitions. One of the primary assumptions is that control signals have a one-to-one mapping with brain regions. That is, every control signal gets delivered to one brain region and one region only. This assumption clashes with the reality of neurostimulation, which affects not only a target site, but also proximal off-target areas. Here, we present a simple extension to account for this reality by modifying the input matrix to include spatial dependencies. In this way, input signals are injected into the network in a spatially diffuse manner. We find that this extension takes advantage of spatial regularities in brain activations and brain network topology, reducing the total energy needed for brain state transitions. We also show that, using a similar approach, we can devise an input strategy for achieving near-optimal control but with much smaller set of effective input signals. This approach generates maps of those sites, which we compare against a set of brain annotations, identifying several maps that correspond closely with the input maps and help establish a stronger neurobiological basis for optimal control.

### Spatially diffuse inputs reduce global control energy

Here, we present evidence that, on a per-transition basis, spatially diffuse inputs reduce control energies needed to go from one brain state to another. This observation suggests that previous studies using the standard “local input” strategy may represent overestimates of control energies. The reduction in energy is likely due to spatial correlations in both brain activation maps and the connectome itself. If two spatially proximal nodes have similar initial/target activation patterns, similar connectivity patterns, and require similar input signals, the local input strategy would nonetheless need to stimulate both nodes independently. In contrast, a single spatially diffuse input signal could, in principle, be used to deliver inputs to both nodes simultaneously. This observation is in line with studies documenting strong spatial autocorrelation in meso-/macro-scale connectomes [65, 66] and functional imaging data [67, 68].

Interestingly, we found that this effect – wherein the control energy was reduced for spatial inputs compared to local inputs – was restricted to a specific range of the diffusivity parameter. For small parameter values, corresponding to exceptionally broad spatial inputs, the local strategy actually outperforms the spatial strategy. This is likely due to the fact that, in this range, the inputs effectively have no specificity in terms of which nodes they impact and therefore has implications for neurostimulation studies [69–72]. Namely, it predicts that a given stimulation technique – e.g. transcranial magnetic stimulation – may be inefficacious in driving the brain into specific states if it cannot achieve stimulation precision below a specific threshold. Here, we can provide an estimate of that threshold based on Fig. 3e. The “crossover” point occurs when *β* ≈ 0.05. At this parameter value, the exponential function decays to half its peak by approximately 15 mm, falling off sharply for longer distances.

### Approximate control with few inputs

One of the challenges associated with optimal control is to identify not only the lowest-energy input signals, but also the fewest *number* of inputs and their locations (control sites). Previous studies have been forced, due to mathematical constraints, to solve for the optimal inputs using a relatively large number of input sites [21, 22]. In fact, due to this limitation, most contemporary applications of optimal control deliver inputs through every brain region [23, 25, 37, 38].

Here, we show that control signals are highly compressible and redundant. That is, when we consider the shape and amplitude of the control signal, we find that many regions receive similar patterns of inputs, suggesting that the total number of independent inputs can be reduced to a much smaller set of relevant “exemplar” input signals (either *via* clustering, as we did here, or through other dimension reduction techniques). Here, we used cluster labels to obtain “input maps,” which highlighted the spatial extent of each cluster and revealed regions that receive similar inputs. Broadly, the maps are distributed across cortex and vary smoothly, reinforcing the “network” perspective of brain organization and function. That is, that brain regions do not act independently; rather, they function collectively as spatially distributed network entities [50] and their inputs, in the context of optimal control and brain stimulation, should be tailored accordingly.

We also show that the input maps exhibit a broad correspondence with other well-established brain maps. On one hand, this is unsurprising – the input maps are closely aligned with the activation patterns of the eleven brain states and many studies have documented statistical associations between maps of brain function and other annotations [73–75]. On the other hand, our findings complement recent work seeking to understand the biological basis of optimal control [38], identifying a series of markers that may play a role in dictating the efficacy of control.

### Future directions

This study presents a number of opportunities for future studies. One such extension involves the decision to model the spatial dependence of the input signals as a decaying exponential; the framework presented here is generic and amenable to many different spatial relationships. For instance, it would be straightforward to model the effect of space as a Gaussian distribution (centered on a given control site) or a “Mexican hat” function that depends on spatial proximity.

The spatial input strategy opens up other unique opportunities for exploring network control. For instance, one could model the effect of a control signal whose input site does not overlap with the centroid of a given parcel/node. That is, we could imagine a control signal originating at a location positioned exactly on the border of two parcels (a scenario that the current optimal control framework would disallow), but under the spatial model we could calculate the effect of that input signal on each node by weighting its effect on either node based on their distance from the input site.

Another important extension of this work involves exploring the efficacy of spatially diffuse control for transitions other than the discrete set of empirically-derived brain states considered here. That is, for other states – e.g. task activations or meta-analytic maps – does the spatial model hold an advantage over the local input strategy? Future work should investigate this question in greater detail.

A final extension includes the exploration of alternative distance metrics for defining the spatial input matrix. Here, we used Euclidean distance between parcel centers of mass to determine the spatial diffusivity of control signals. However, if we imagine that the diffusivity reflects volume conduction, then our distance metric should take into account cortical folding patterns. Accordingly, a more appropriate distance metric would be the geodesic distance between regions along the cortical surface. We leave this exploration for a future study.

### Limitations

This study has a number of limitations. Here, we focus on large-scale human connectome data estimated from tractography. The procedure for reconstructing connectome data suffers from well-known biases [56, 76, 77]. Because control energies – estimated under both the “local” and “spatial” strategies – depend on the topology of the network, any error in the estimate of the connectome will propagate to errors in the energy estimate.

An additional limitation concerns the exclusion of subcortical regions of interest from this analysis. The decision to focus on only neocortex was motivated practically. Namely, most standard subcortical parcellations are developed in isolation and are integrated “as is” into existing pipelines for parcellating neocortex. Oftentimes, this results in parcels whose volumes differ (sometimes dramatically) from those of cortical parcels, which can inadvertently inflate estimates of connectivity. For instance, combining small cortical parcels with large subcortical parcels can give the impression that subcortical regions are connected to virtually every cortical ROI. While recent advances have helped remedy this [78], it remains an open issue. Future work should be focused on jointly parcellating and modeling cortical/subcortical/cerebellar connectomes.

## MATERIALS AND METHODS

### Datasets

In this study we analyzed two separate MRI datasets. We refer to the primary dataset, reported on in the main text, as the “Lausanne dataset,” and the replication dataset as the “Human Connectome Project dataset.”

#### Lausanne dataset

In this study we examined optimal control in MRI-defined connectomes. We carried out these comparisons using diffusion spectrum MRI data parcellated networks at a single organizational scale (*N* = 1000 cortical nodes). Here, we describe those processing steps in greater detail.

Informed written consent in accordance with institutional guidelines (protocol approved by the Ethics Committee of Clinical Research of the Faculty of Biology and Medicine, University of Lausanne, Switzerland, #82/14, #382/11, #26.4.2005) was obtained for all subjects. Data provided are fully anonymized. A total of 70 healthy participants (age 28.8 *±* 9.1 years, 27 females) were scanned in a 3-Tesla MRI scanner (Trio, Siemens Medical, Germany) using a 32-channel head-coil. The session protocol was comprised of (1) a magnetization-prepared rapid acquisition gradient echo (MPRAGE) sequence sensitive to white/gray matter contrast (1-mm in-plane resolution, 1.2-mm slice thickness), (2) a DSI sequence (128 diffusion-weighted volumes and a single b0 volume, maximum b-value 8,000 s/mm2, 2.2×2.2×3.0 mm voxel size), and (3) a gradient echo EPI sequence sensitive to BOLD contrast (3.3-mm in-plane resolution and slice thickness with a 0.3-mm gap, TR 1,920 ms, resulting in 280 images per participant). During the fMRI scan, participants were not engaged in any overt task, and the scan was treated as eyes-open resting-state fMRI (rs-fMRI).

Initial signal processing of all MPRAGE, DSI, and rs-fMRI data was performed using the Connectome Mapper pipeline [44]. Gray and white matter were segmented from the MPRAGE volume using freesurfer [79] and parcellated into 83 cortical and subcortical areas. The parcels were then further subdivided into 129 (114 cortical), 234 (219 cortical), 463 (448 cortical) and 1015 (1000 cortical) approximately equally sized parcels according to the Lausanne anatomical atlas following the method proposed by [80].

DSI data were reconstructed following the protocol described by [81], allowing us to estimate multiple diffusion directions per voxel. The diffusion probability density function was reconstructed as the discrete 3D Fourier transform of the signal modulus. The orientation distribution function (ODF) was calculated as the radial summation of the normalized 3D probability distribution function. Thus, the ODF is defined on a discrete sphere and captures the diffusion intensity in every direction.

Structural connectivity matrices were estimated for individual participants using deterministic streamline tractography on reconstructed DSI data, initiating 32 streamline propagations per diffusion direction, per white matter voxel [82]. Within each voxel, the starting points were spatially random. For each starting point, a fiber streamline was grown in two opposite directions with a fixed step of 1 mm. Once the fiber entered a new voxel, the fiber growth continued along the ODF maximum direction that produces the least curvature for the fiber (i.e., was most similar to the trajectory of the fiber to that point). Fibers were stopped if the change in direction was greater than 60 degrees/mm. The process was complete when both ends of the fiber left the white matter mask. Structural connectivity between pairs of regions was measured in terms of fiber density, defined as the number of streamlines between the two regions, normalized by the average length of the streamlines and average surface area of the two regions [83]. The goal of this normalization was to compensate for the bias toward longer fibers inherent in the tractography procedure, as well as differences in region size.

Functional data were pre-processed using routines designed to facilitate subsequent network exploration [84, 85]. fMRI volumes were corrected for physiological variables, including regression of white matter, cerebrospinal fluid, as well as motion (three translations and three rotations, estimated by rigid body co-registration). BOLD time series were then subjected to a lowpass filter (temporal Gaussian filter with full width half maximum equal to 1.92 s). The first four time points were excluded from subsequent analysis to allow the time series to stabilize. Motion “scrubbing” was performed as described by [85].

Rather than analyze data from individual participants, we constructed a group-representative connectome. The typical procedure for doing is to apply a consensus threshold across edges, retaining only those edges that appear in a fixed fraction of participants. This fraction is generally uniform across all edges. However, this procedure generates consensus networks with fewer long-distance connections and more short-range connections than the typical subject. To circumvent these issues, we used a distance-dependent consensus algorithm for generating the group-representative networks [86]. All analyses were carried out using this network.

#### Human Connectome Project dataset

The Human Connectome Project (HCP) dataset [43] consists of structural magnetic resonance imaging (T1w), resting state functional magnetic resonance imaging (rsfMRI) data, and diffusion MRI (dMRI) data from young adult subjects. The dataset spans two collection sites, at Washington University in St. Louis on a 3T MRI machine. For the present study, the MRI data consist of a subset of 100 unrelated adults (“100 Unrelated Subjects” published by the Human Connectome Project). The study was approved by the Washington University Institutional Review Board and informed consent was obtained from all subjects. HCP 3T data were quality controlled based on motion summary statistics and visual inspection. After exclusion of four high motion subjects and one subject due to software error, the final HCP 3T subset consisted of 95 subjects the final subset utilized included 95 subjects (56% female, mean age = 29.29 *±* 3.66, age range = 22-36).

A comprehensive description of the HCP imaging parameters and image prepocessing can be found in [87] and in HCP’s online documentation (https://www.humanconnectome.org/study/hcp-young-adult/document/1200-subjects-data-release). For all HCP subjects, T1w were collected on a 3T Siemens Connectome Skyra scanner with a 32-channel head coil. Subjects underwent two T1-weighted structural scans, which were averaged for each subject (TR = 2400 ms, TE = 2.14 ms, flip angle = 8^*°*^, 0.7 mm isotropic voxel resolution). For all fMRI data collected, 4 gradient-echo planar imaging sequences were collected for each subject: two runs were acquired with left-to-right phase encoding direction and two runs were acquired with right-to-left phase encoding direction.

HCP 3T fMRI was collected on a 3T Siemens Connectome Skyra with a 32-channel head coil. Each resting state run duration was 14:33 min, with eyes open and instructions to fixate on a cross (TR = 720 ms, TE = 33.1 ms, flip angle = 52^*°*^, 2 mm isotropic voxel resolution, multiband factor = 8).

Structural and functional HCP images were minimally preprocessed according to the description provided in [87]. Briefly, T1w images were aligned to MNI space before undergoing FreeSurfer’s (version 5.3) cortical reconstruction workflow. fMRI images were corrected for gradient distortion, susceptibility distortion, and motion, and then aligned to the corresponding T1w with one spline interpolation step. This volume was further corrected for intensity bias and normalized to a mean of 10000. This volume was then projected to the 2mm *32k_fs_LR* mesh, excluding outliers, and aligned to a common space using a multi-modal surface registration [88]. The resultant cifti file for each HCP subject used in this study followed the file naming pattern: `*_Atlas_MSMAll_hp2000_clean.dtseries.nii`.

All resting state fMRI images were nuisance regressed in the same manner. Each minimally preprocessed fMRI was linearly detrended, band-pass filtered (0.008-0.08 Hz), confound regressed and standardized using Nilearn’s `signal.clean` function, which removes confounds orthogonally to the temporal filters. We used the following confound regression strategy, termed “aCompCor”. Briefly, this approach included six motion estimates, derivatives of these previous six regressors, and squares of these 12 terms, in addition to five anatomical CompCor components [89]; aCompCor is a non-GSR strategy. Following these preprocessing operations, the mean signal was taken at each time frame for each node, forming nodal time series. Cortical nodes were defined by the Schaefer 400 cortical parcellation [90] in the *32k_fs_LR* surface space.

dMRI images were normalized to the mean b0 image, corrected for EPI, eddy current, and gradient non-linearity distortions, and motion, and aligned to subject anatomical space using a boundary-based registration [91]. In addition to HCP’s minimal preprocessing, diffusion images were corrected for intensity non-uniformity with `N4BiasFieldCorrection` [92]. FSL’s `dtifit` was used to obtain scalar maps of fractional anisotropy, mean diffusivity, and mean kurtosis. The Dipy toolbox (version 1.1) [93] was used to fit a multi-shell multi-tissue constrained spherical deconvolution [94] to the diffusion data with a spherical harmonics order of 8, using tissue maps estimated with FSL’s `fast` [95]. Tractography was performed using Dipy’s `Local Tracking` module [93]. Multiple instances of probabilistic tractography were run per subject [96], varying the step size and maximum turning angle of the algorithm. Tractography was run at step sizes of 0.25 mm, 0.4 mm, 0.5 mm, 0.6 mm, and 0.75 mm with the maximum turning angle set to 20^*°*^. Additionally, tractography was run at maximum turning angles of 10^*°*^, 16^*°*^, 24^*°*^, and 30^*°*^ with the step size set to 0.5 mm. For each instance of tractography, streamlines were randomly seeded three times within each voxel of a white matter mask, retained if longer than 10 mm and with valid endpoints, following Dipy’s implementation of anatomically constrained tractography [97], and errant streamlines were filtered based on the cluster confidence index [98].

For each tractography instance, streamline counts were normalized by dividing the count between nodes by the geometric average volume of the nodes. Since tractography was run nine times per subject, edge values were collapsed across runs. To do this, the weighted mean was taken with weights based on the proportion of total streamlines at that edge. This amounts to calculating the expected value, where probabilities are based on the proportion of total edge weight across tracotgraphy instances. This operation biases edge weights towards larger values, which reflect tractography instances better parameterized to estimate the geometry of each connection.

As with the Lausanne dataset, we focused on a group-representative network with distance-dependent consensus binning [86].

### Brain state detection

We z-scored parcel (nodal/regional) time series for each participant and calculated at each frame the root mean square activity. That is, if *x*_*i*_(*t*) corresponds to the activity of region *i* at time *t*, we calculated 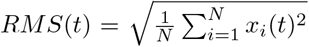 . We then identified peaks in the *RMS* time series as any frame whose preceding and following frame had smaller *RMS*, collected the corresponding peak activation patterns across participants, and calculated the *N*_*peaks*_ × *N*_*peaks*_ concordance matrix [99].

The concordance between between two vectors *x* and *y* is calculated as:

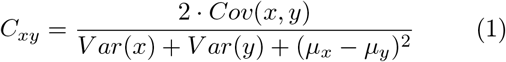

where *μ*_*x*_ and *V ar*(*x*) are the sample mean and variance of *x*, and *Cov*(*x, y*) is the covariance between *x* and *y*. Intuitively, if the variances and means of *x* and *y* are identical, then this expression evaluates to the correlation coefficient, *r*_*xy*_. However, if those statistics difsfer, then *C*_*xy*_ *< r*_*xy*_. That is, *C*_*xy*_ is a measure of statistical similarity that penalizes for differences in mean and variance. The concordance metric has been used in previous studies for brain state detection [46].

We then use modularity maximization to identify clusters in the concordance matrix. Briefly, modularity maximization is a heuristic for discovering communities (clusters) in graphs [45]. Intuitively, it defines communities as groups of nodes whose density of connectivity to one another maximally exceeds that of a null model. This intuition can be formalized with quality function:

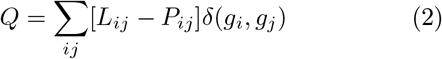

where *L*_*ij*_, in this case, represents the concordance between peak activation maps *i* and *j, δ*(*·, ·*) is the Kronecker delta function, which evaluates to 1 when its inputs are identical and 0 otherwise, and *g*_*i*_ is the community to which activation map *i* is assigned.

It is up to the user how to define *P*_*ij*_ [100], the expected concordance between maps *i* and *j*. Here, we define *P*_*ij*_ = *mean*(**W**) + *stdv*(**W**). That is, we set the expected concordance to be the mean concordance between all pairs of activation maps plus one standard deviation.

We use the Louvain algorithm to optimize *Q* with 1000 random restarts. This procedure yields 1000 dissimilar estimates of communities, which we combine to generate a single set of consensus communities [101]. Briefly, the consensus community algorithm works by calculating for every pair of peak activation maps the fraction of the 1000 Louvain runs in which they were assigned to the same community. It also calculates the expected co-assignment probability (by randomly permuting the detected community labels). This allows us to compare the observed and expected coassignment probabilities, setting up a consensus modularity function, which we optimize using the same Louvain algorithm. Clustering the consensus matrix reduces variability across the detected communities, oftentimes yielding a single solution across the 1000 runs. If that is the case, then the algorithm has discovered the “consensus clusters”. If variability remains, the consensus algorithm is applied again to the new estimates of communities. These steps – generating estimates of communities and calculating the coassignment matrix – are repeated until the estimated clusters are all identical.

Each community is composed of activation maps whose mean pairwise concordance exceeded the expected concordance. We calculated community centroids as the midpoint across all maps assigned to a given community. We also calculated, for each community, the fraction of the 70 participants with at least one activation assigned to that community. We retained those communities in which at least 50% of participants were represented.

Here we clustered whole-brain activity patterns corresponding to peaks in the activity time series. Alternatively, one could cluster activity patterns at every frame. However, there are at least two reasons why this may be sub-optimal. First, it would result in orders of magnitude more patterns, drastically lengthening the runtime of the clustering algorithm and limiting its utility for larger datasets. Second, clustering the complete set of activity patterns would likely not result in the detection of novel brain states; the strong autocorrelation observed in the fMRI BOLD data ensures that temporally proximal frames are highly similar [102]. Thus, the peak sampling approach can be viewed as a means of downsampling the fMRI data and reducing biases that arise from clustering highly autocorrelated samples.

### Optimal control

We consider the linear dynamical system:

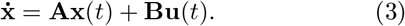

Here, **x**(*t*) = [*x*_1_(*t*), …, *x*_*N*_ (*t*)] is the brain state at time *t*; **A** ∈ *ℝ*^*N*×*N*^ is the weighted (and in this case) symmetric connectivity matrix whose element *A*_*ij*_ denotes the presence/absence of the connection between nodes *i* and *j*; **u**(*t*) is the set of control inputs, with one input per control point; **B** is the input matrix, which maps control inputs to network nodes.

Note that here we normalize the connectome so that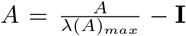, where *λ*(*A*)_*max*_ is the largest eigen-value of *A* and **I** is the identity matrix. This normalization step ensures that all of **A** eigenvalues are negative, prohibiting explosive growth.

Taken together, this equation implies that, in the absence of any input signals the system evolves passively according to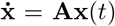. However, if the control set is not empty then the evolution of nodes’ states will depend on the control signal **u** and where it gets injected into the system, which is specified by the coupling matrix, **B**.

Optimal control is an engineering framework that seeks to drive a networked dynamical system from an initial brain state, **x**_0_ = **x**(*t* = 0) into a desired target state, **x**_*T*_ = **x**(*t* = *T*), where the time *t* = *T* is referred to as the “control horizon.”

The optimal control framework seeks to minimize the quantity:

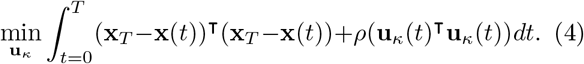

We can think of this minimization problem as being composed of two terms. The first term, (**x**_*T*_ − **x**(*t*))^⊺^(**x**_*T*_ − **x**(*t*)), corresponds to the distance of the brain’s state at time *t* from its target state. The second term, **u**_*κ*_(*t*)^⊺^**u**_*κ*_(*t*), corresponds to amplitude of the control signals. The parameters *T* and *ρ* refer to the control horizon and the balance between the two terms to be minimized, respectively. Here, we follow previous studies [21, 22] and set *T* = 1 and *ρ* = 100. The variable *κ* denotes the set of input sites. Here, we consider the case where all brain regions receive control signals, i.e. *κ* = *{*1, …, *N }*.

To identify the optimal inputs, **u**_*κ*_, we define the Hamiltonian:

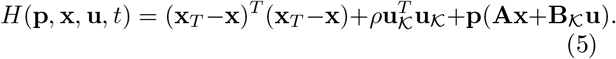

From the Pontryagin minimization principle, if 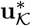 is an optimal solution to the minimization problem with corresponding trajectory, **x**^*^, then there exists **p**^*^ such that:

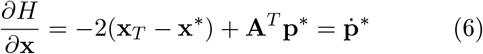

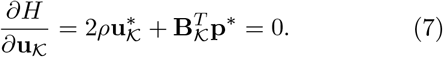

This set of equations reduces to:

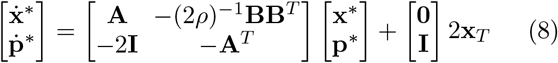

If we denote:

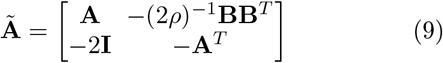

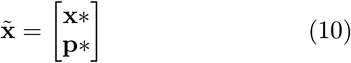

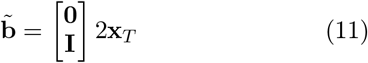

we can then write the reduced equation as:

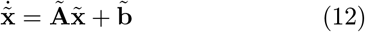

which we can solve as:

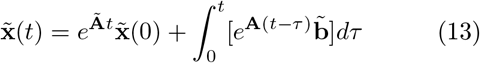

or, alternatively

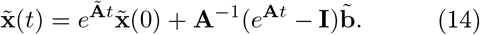

Then, substituting *t* = *T*, we arrive at:

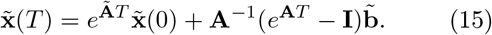

Let

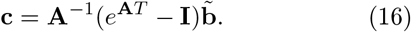

We can then write:

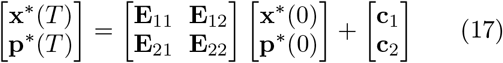

Rewriting this, we get:

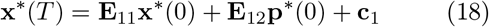

which can be rearranged to write:

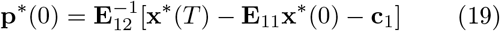

Given **p**^*^(0) and **x**_0_, we can then integrate 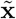forward, thereby obtaining **x**_*T*_ from which we subsequently obtain the optimal inputs, 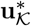.

Note, in this derivation of the optimal inputs, we need only specify the free parameter *ρ* and the boundary conditions, namely **x**_*T*_ and **x**_0_. Collectively, these variables determine the value of **p**^*^(0). It is also worth noting that while some of the variables have clear physical interpretations (e.g. (**x**_*T*_ − **x**)^*T*^ (**x**_*T*_ − **x**) is the distance from the target state and 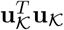 is related to the total energy) other variables do not. The additional variables that appear in this section are a consequence of the optimization technique that we used and come from the technique of Lagrange multipliers [103].

### A spatially diffuse input matrix

In its “classical” form, the input matrix, **B**, is composed of indicator column vectors. The elements of the *i*th vector are all equal to zero except that of row *i*, whose value is set equal to 1; this ensures that the corresponding input signal is delivered only to node *i* and nowhere else. If all nodes are treated as control points, then **B** = **I**, where **I** ∈ ℝ^*N*×*N*^ is the identity matrix. We denote the “local” input matrix as **B**^*local*^.

Although convenient, strictly local inputs are not compatible with existing methodologies for perturbing large-scale brain systems nor do they agree with dominant theories of brain function. Specifically, injecting an input signal into a region *i* such that the effect of the perturbation is circumscribed to that region of interest, alone, is not plausible. Rather, stimulation to region *i* is “felt” by regions spatially proximal to that of *i*. Additionally, it is well established that brain regions (nodes) behave in concert with one another as part of distributed brain-wide networks. This suggests that the “local” input strategy, which assumes independence of the input signals, is not plausible.

Instead, we can incorporate some realism by modeling the effect of space on input signals. Specifically, we model this effect as a negative exponential [68, 104]. That is, region *j* receives the following contribution from the input signal centered on *i*:

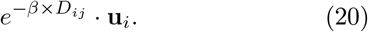

Here, *D*_*ij*_ is the Euclidean distance between the centers of mass for regions *i* and *j*. The parameter *β* determines the rate at which the spatial effect diminishes. As *β* → ∞, the value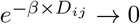.

More generally, we can think of the above expression as defining the elements of a spatially-diffuse input matrix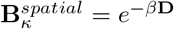.

### Approximations of optimal control

In the main text we described an algorithm for approximating optimal control using a much smaller set of input signals. The algorithm for doing so proceeds as follows. First, using every node as an input site, we obtain the optimal input signals. We then use the *k*-means clustering algorithm (varying the number of clusters from *k* = 2 to *k* = 40; Euclidean distance) to partition input signals based on their temporal similarity. We then estimate the mean input signal for each cluster, *c*:

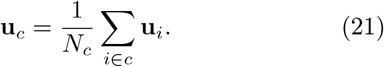

Next, we calculate a composite input matrix using a similar strategy:

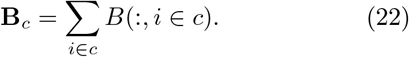

Here, *B*(:, *i* ∈ *c*) refers to the *i*th column of the input matrix.

Given the low-dimensional input matrix, **B**_*c*_, and control signals, **u**_*c*_, and subject to the boundary constraint such that **x**(*t* = 0) = **x**_0_, we can integrate this system to obtain an estimate of the target brain state **x**^*′*^(*t* = *T*). In general, because the input signals are inexact (due to the clustering procedure), **x**^*′*^(*t* = *T*)*≠* **x**_*T*_ . We can then calculate the discrepancy between the approximation of the target state with the true target state as:

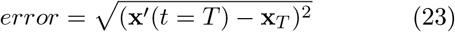

### Brain annotations

A curated set of brain maps were fetched from the *neuromaps* toolbox [59] and parcellated into 1000 equally-sized parcels using the same atlas used to parcellate the connectivity data [80]. More information on each individual brain map is available in the toolbox’s repository (https://github.com/netneurolab/neuromaps). Briefly, the annotations included 27 neurotransmitter receptor and transporter PET tracer density maps [105] from nine different neurotransmitter systems including serotonin [106, 107], dopamine [108–112], norepinephrine [113, 114], histamine [115], acetylcholine [105, 116–119], cannabinoid [120, 121], opioids [122–124], glutamate [125, 126] and GABA [127]. Receptor/transporter with more than one mean density map were combined to obtain a total of 19 density maps for individual neurotransmitter receptors and transporters. Listed in italics are the labels used in the text for each: *5HT1a, 5HT1b, 5TH2a, 5HT4, 5HT6, 5HTT, a4b2, CB1, D1, D2, DAT,GABAa, H3, KOR, M1, mGluR5, MOR, NAT, VAChT*. The annotations also included 10 maps of diffusion map embedding gradients of group-averaged functional connectivity (*fcgradient01* to *fcgradient10*) [64], 5 maps of metabolic measures including cerebral blood flow (*cbf* and *cbfmean*) and volume (*cbv*), oxygen metabolism (*cmro2*), and glucose metabolism (*cmrglu*) [128, 129], 6 maps of MEG frequency power distribution (*megalpha, megbeta, megdelta, meggamma1, meggamma2, megtheta*) and a map of MEG-derived intrinsic time scale (*megtimescale*) [43, 130]. Also included are 2 maps of morphometric MRI measures: T1w/T2w ratio acting as a proxy for myelin (*myelinmap*) and cortical thickness (*thickness*) [131], a map of cross-species functional homology (*Fchomology*) [132], a map of evolutionary cortical expansion (*evoexp*) [132], the sensory-association mean rank axis (*SAaxis*) [60], the principal component of neurosynth terms in the cognitive atlas (*cogPC1*) [62, 63], the principal component of gene expression (*genePC1*) [133, 134], a map of intersubject variability of resting-state functional connectivity (*intersubjvar*) [135], a map of synapse density (*SV2A*) [136] and 3 maps of cortical areal scaling during development [137], which were averaged and combined into a single annotation (*scaling*).

## ACKNOWLEDGEMENTS

RFB acknowledges support from the National Science Foundation (award number 2023985) and National Institute on Aging (project number 5R01AG075044-02). LP was supported by the National Institute Of Mental Health of the National Institutes of Health under Award Number R00MH127296.

---

**Figure S1.**
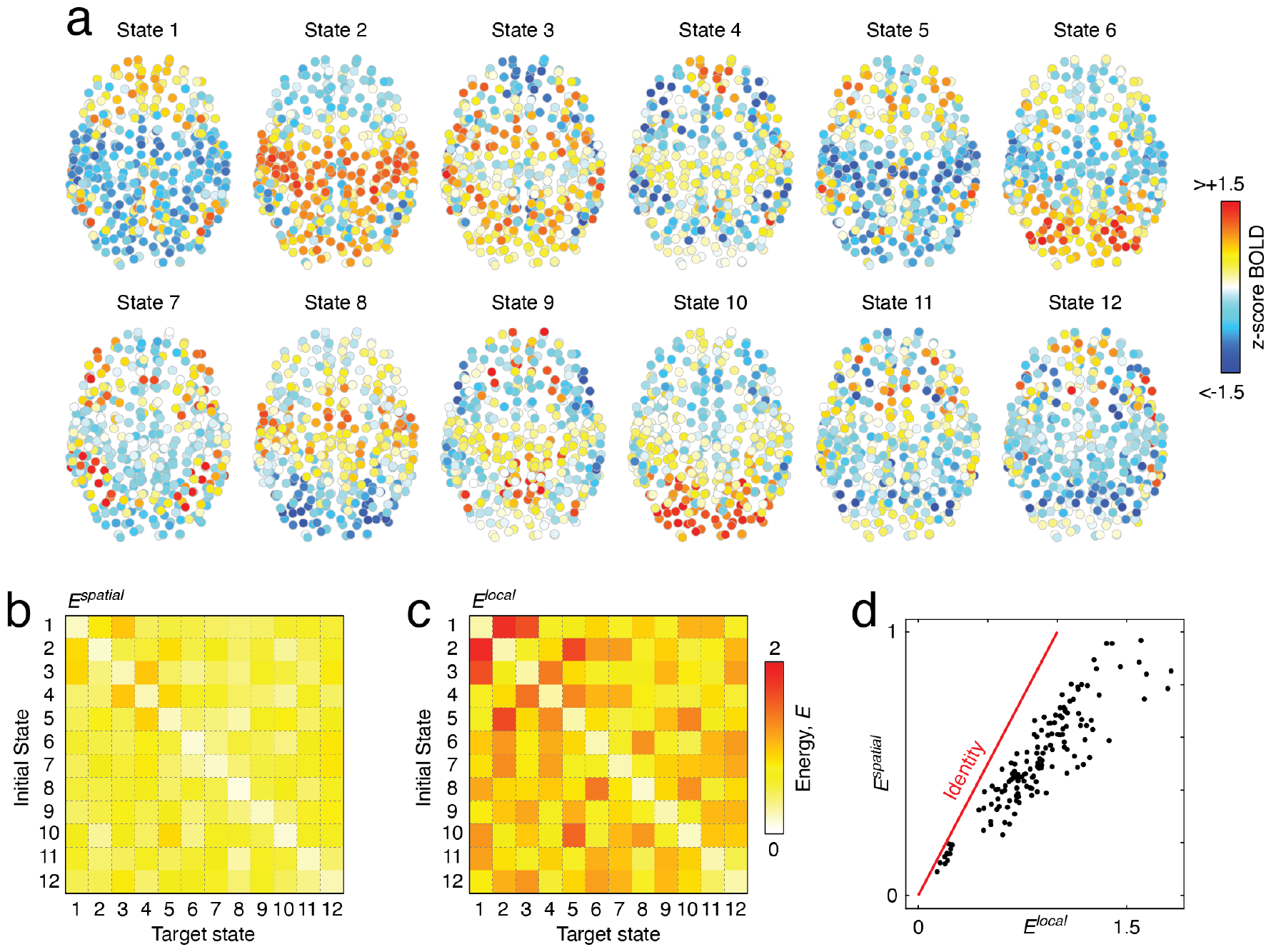
Independent replication of main finding. In the main text we showed that spatially diffuse inputs reduce the control energy needed to transition between a set of 11 empirically defined brain states compared to the traditional “local” input strategy. Here, we replicate this finding using a larger and independently acquired and processed dataset. Specifically, we used resting-state and diffusion MRI data from the Human Connectome Project. In general, we followed an analysis pipeline identical to the one described in the main text. We used peak sampling to obtain 6241 peaks in the resting-state activity (`REST1_LR`) from 95 unrelated participants with complete datasets that also met minimum data quality standards. (*a*) After clustering the peak activity patterns, we identified 12 brain states that appeared in at least 50% of the participants. We show the topography of those states here. Panels *b* and *c* show control energies for all pairs of brain state transitions. (*d*) Scatter plot of energies. For all 144 transitions, the spatial input strategy yielded lower energies than the local strategy.

**Figure S2.**
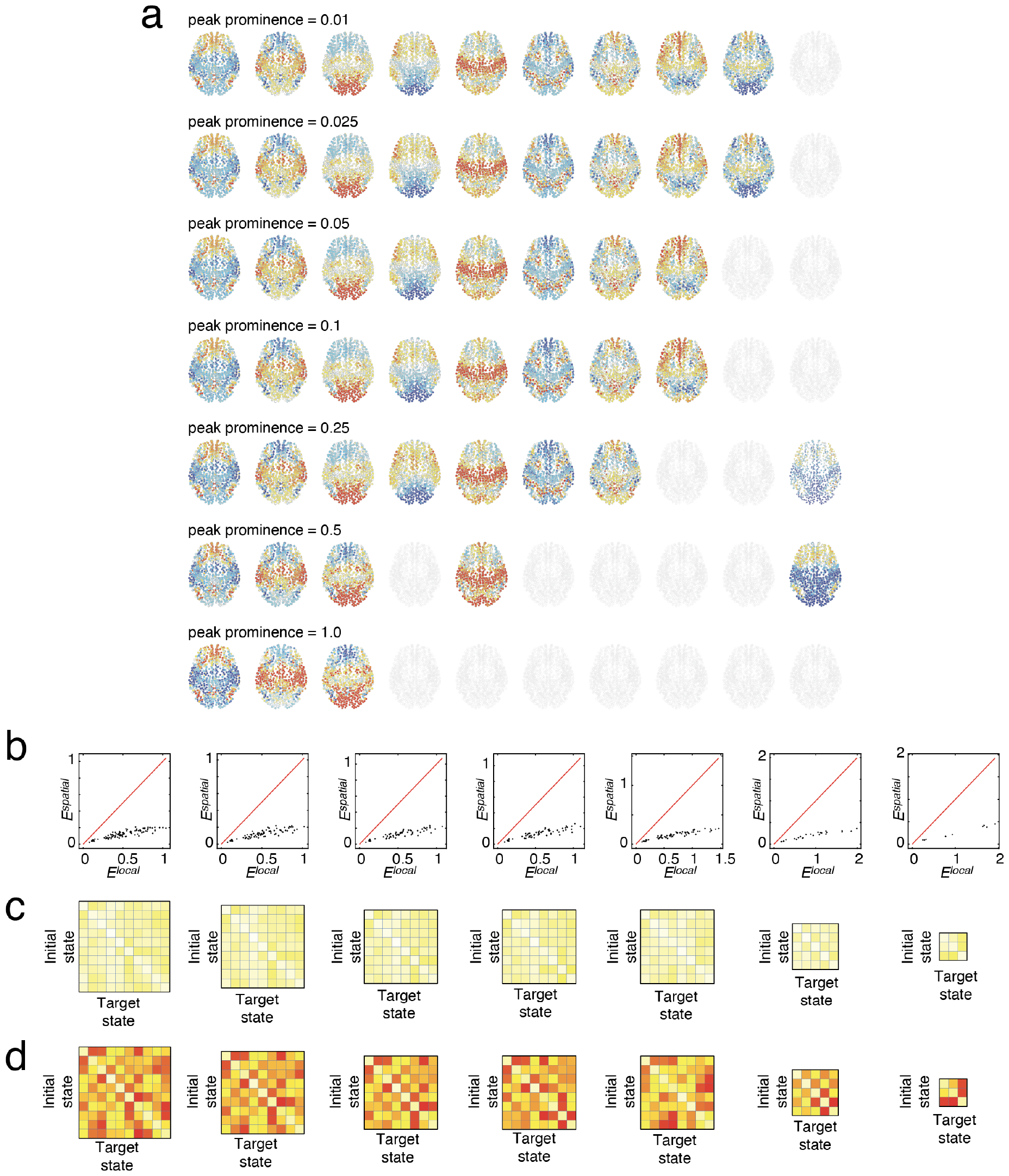
Alternative state definitions. In the main text we showed that spatially diffuse inputs reduce the control energy needed to transition between a set of 11 empirically defined brain states compared to the traditional “local” input strategy. The brain states we defined by sampling and clustering whole-brain activation patterns from peaks in the global signal. However, some of these peaks may reflect “noise” – i.e. though technically peaks, the peak may not be prominent. Here, we explore the effect of alternative brain state definitions in which we retain only the most prominent peaks by adjusting the ‘prominence” parameter (the drop on both sides of the peak) from 0.01, 0.025, 0.05, 0.1, 0.25, 0.5, and 1. (*a*) Cluster centroids at each prominence value. Note that, as in the main text, we retain only those clusters in which at least 50% of the participants were represented. Cluster centroids have been aligned visually so that columns reflect, approximately, the same clusters across parameter values. Treating the clusters as estimates of “brain states”, we calculated all pairwise transitions under both the spatial and local input strategies. (*b*) Scatter plot comparing energies under spatial and local models. As in the main text, on a per-transition basis, the spatial energy was consistently lower than the local energy. Panels *c* and *d* show energy values for the spatial input strategy (panel *c*) and local strategy (panel *d*).

**Figure S3.**
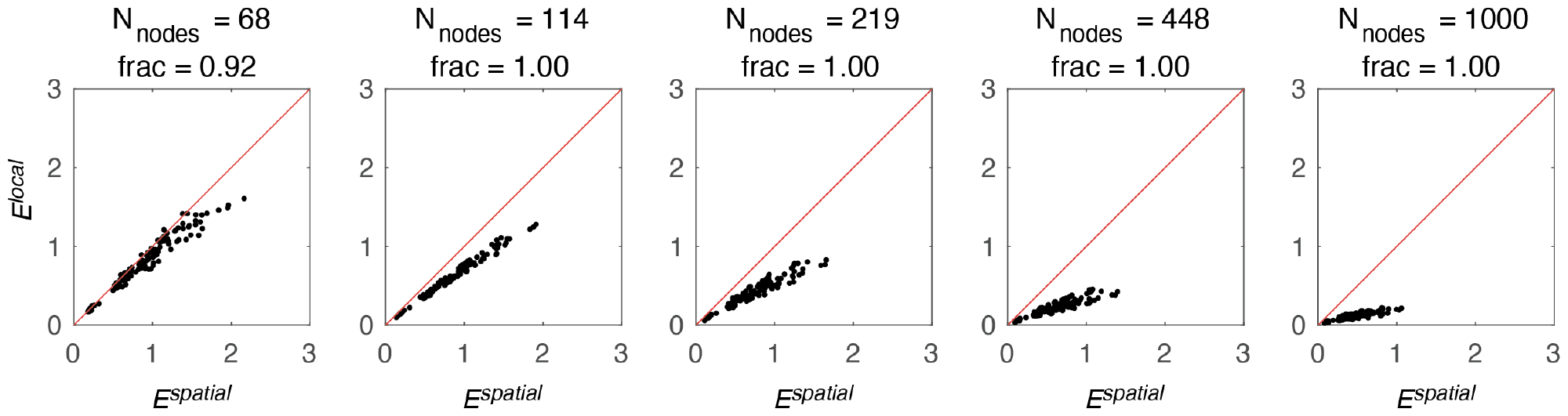
Alternative parcel definitions. The “Lausanne” parcellation is multiscale. The five scales contain 68, 114, 219, 448, and 1000 cortical parcels. We repeated our main analysis using connectomes and brain states defined at these other scales. In general, we found that our results replicated. Here, the *x* and *y* axes correspond to the energy for each transition under the spatial and local input strategies, respectively. In the title of each plot, “frac” refers to the fraction of transitions in which the spatial model yielded a smaller energy compared to the local model. That is, a value of 1.00 means that all 121 transitions resulted in lower energy under the spatial model.

**Figure S4.**
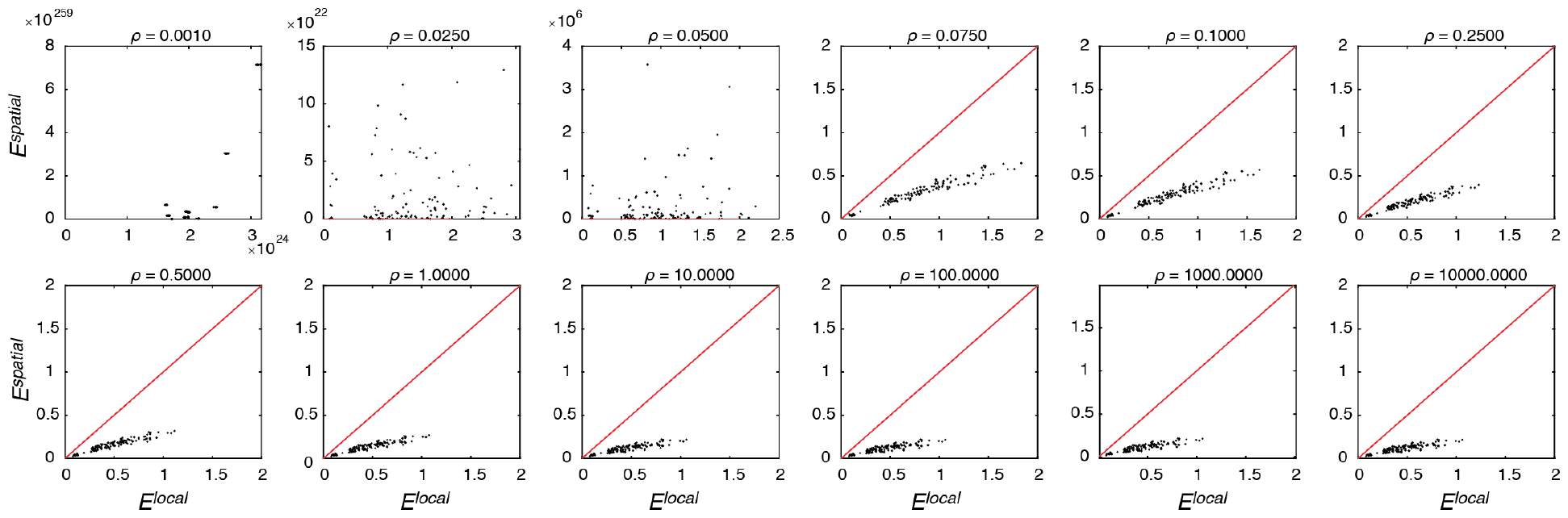
Effect of varying *ρ*. In the optimal control framework, the parameter *ρ* controls the balance of the two terms in the optimization: larger values of *ρ* upweight the importance of the control energy relative to the distance from the reference state. Here, we show that, across wide ranges of *ρ* (spanning roughly six decades, from *ρ* = 10^−1^ to at least *ρ* = 10^4^), the spatial input strategy results in reduced control energy compared to the local input strategy. At extremely small values of *ρ*, we find numerical instabilities, resulting in energies on the order of 10^100^, and inaccurate estimates of the optimal control signal, making comparisons between the two input strategies difficult.

**Figure S5.**
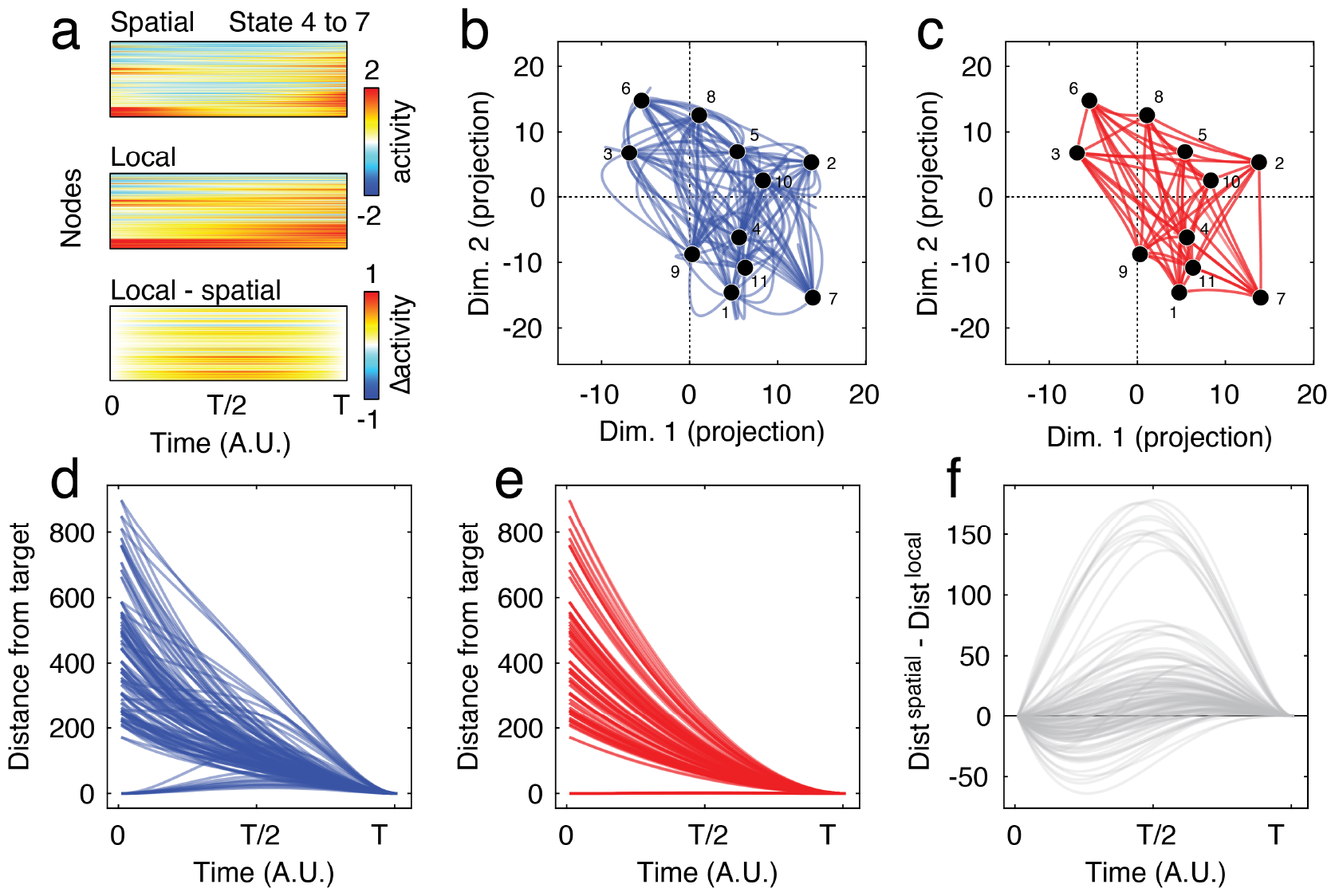
Brain state trajectories under spatial *versus* local control strategies. (*a*) Heatmaps showing nodes’ states across an example control task (a transition from state 4 into state 7). Trajectories were estimated using a diffusivity parameter of *β* = 0.15. The top plot shows node state changes under the spatial input strategy, the middle shows local, and the bottom plot local minus spatial. In the main text, we showed that state trajectories were less direct under the spatial input strategy. That is, the trajectory was allowed to move further from the target state than compared to the local input strategy. To visualize this, we projected brain states onto two random *N* × 1 vectors whose elements were sampled from a normal distribution with zero mean and unit variance. In panel *b*, we show low-dimensional representations of the transitions between all pairs of states under the spatial input strategy. Note that the trajectories tend to be curved. In panel *c* we show an analogous plot for transition under the local strategy. Note that, in this case, the trajectories tend to be straight lines. This effect is evident when we plot the distance from target for each transition as a function of time. Panels *d* and *e* show distances for all 121 transitions (spatial and local control strategies, respectively), while panel *f* shows the differences in distances. Positive/negative values indicate that the trajectory under the spatial/local input strategy was “loopier.”

**Figure S6.**
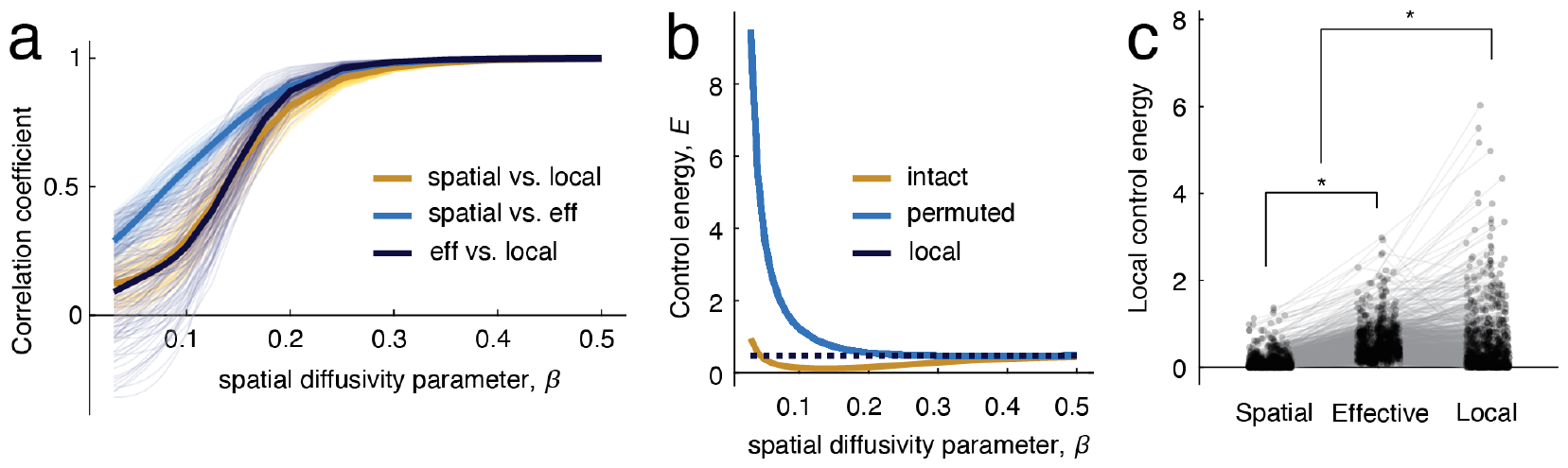
Supplementary analyses. (*a*) Similarity of control energy vectors as a function of the diffusivity parameter, *β*. Each curve corresponds to one of the possible 121 state transitions. (*b*) In the main text we show that the spatial input strategy leads to a reduction in the control energy. We hypothesized that this is because it, effectively, takes advantage of spatial correlations in structural connectivity and brain activations (states). That is, because nearby nodes tend to have similar state values and similar connectivity patterns, weaker but spatially diffuse inputs can have the same effect as the strong but spatially localized inputs. To test this, we simply randomly permuted the rows/columns of the input matrix, thereby destroying any spatial relationships but otherwise preserving exactly its total weight and global structure. We found that when we destroy spatial relationships the control energy increases dramatically. The gold curve is identical to the curve shown in Fig. 3f. The dark blue curve is the energy after destroying spatial structure. (*c*) Regional energies under the spatial input strategy, after calculating the effective energy under the spatial model, and under the local model. We find significant differences between all three distributions.

**Figure S7.**
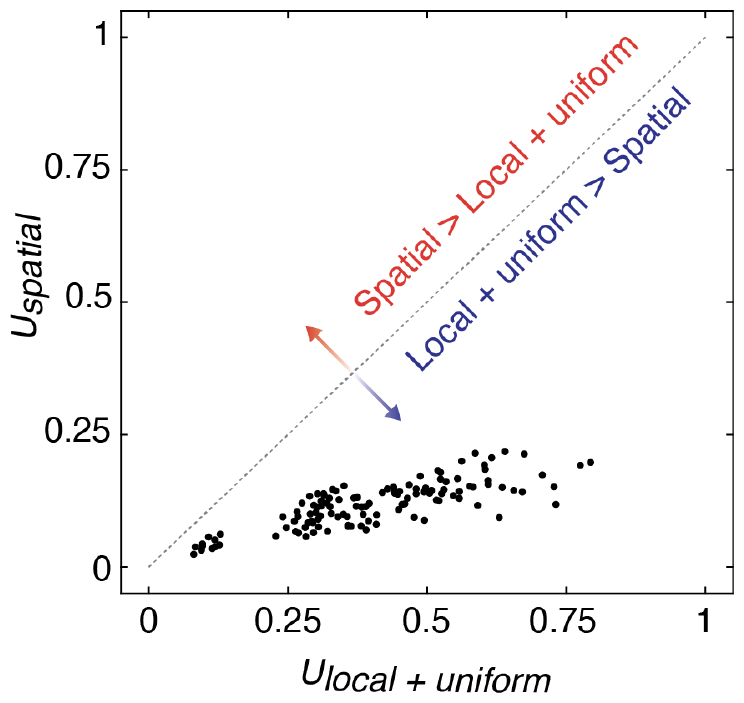
Control energies under null model. The spatial model effectively adds extra nonzero elements to the input matrix. To ensure that the reduction in control energy under the spatial model is not driven exclusively by the added weight, we compared the control energy for all 121 transitions against the energies obtained under the following null model. For column *i*, we calculate 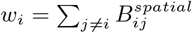 . The value of *w*_*i*_ varies from node to node and corresponds to the additional weight added to column *i* under the spatial model. Next, we initialize a null input matrix as the identity matrix, i.e. *B*_*ii*_ = 1 for all *i*. We then set all off-diagonal elements, *B*_*ij*_ with *i≠ j*, equal to 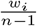. That is, we uniformly distribute the added weight across all non-identity elements within each column. This resulting input matrix has total weight equal to that of the spatial model, its column sums are identical to that of the spatial model, and it preserves the identity line (a feature that both the spatial and local models both exhibit). Next, using this matrix, we calculated the control energy associated with each of the 121 transitions, comparing it to the corresponding energy under the spatial model. We found that the energy under the uniform model was greater than that of the spatial model for every transition. This observation suggests that the observed reduction in energy under the spatial model is not driven solely by the weight added to the input matrix.

**Figure S8.**
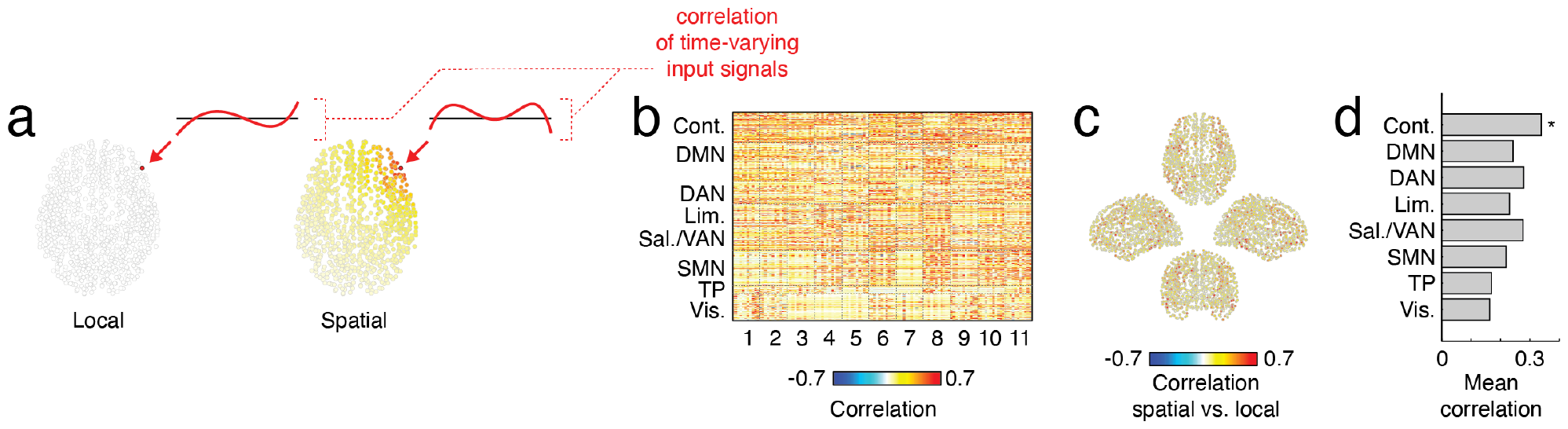
Correlated input signals. In the main text we described a procedure in which we calculated the correlation between regional input signals under the spatial and local input strategies. Here, we elaborate on that procedure. (*a*) Schematic showing input signals being delivered to the same node under different input strategies. Each element in the heatmap shown in *b* corresponds to a correlation coefficient. Rows correspond to nodes and columns correspond to transitions. The columns are ordered by target state. In panel *c* we show, in anatomical space, the mean correlation projected into anatomical space. In panel *d* we show mean correlation at the level of brain systems. The correlation in the control network was the only system whose mean correlation exceeded chance (spin tests).

**Figure S9.**
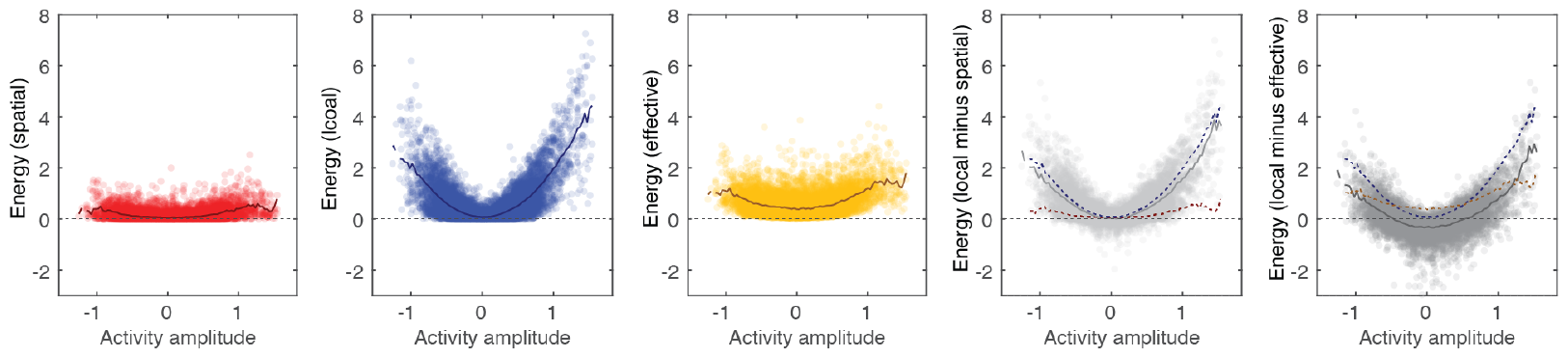
Regions that drive differences between spatial and local input strategies. In the main text we showed that the spatial input strategy reduced control energies compared to the local input strategy. Here, we show that these global reductions are driven by regions whose activity in the target state is extreme (highly active, positively or negatively). The left-most plot (red) depicts regional energies for all control tasks as a function of their activity. The next two plots are analogous to the first; they show (in blue and yellow) regional energies under the local strategy and their effective energy under the spatial strategy. The next panel shows the difference in energy – local minus spatial. It shows that the biggest differences are at the extremes; regions with high activity require proportionally less input under the spatial strategy than under the local strategy. Regions with low levels of activity are comparable between the two input strategies. The right-most panel is analogous to the previous, but shows the difference between local and effective energies. Note that, again, the energy of regions with extreme levels of activity correspond to the biggest gap. However, regions with relatively low levels of activity now receive slightly more energetic inputs compared to the local control strategy.

**Figure S10.**
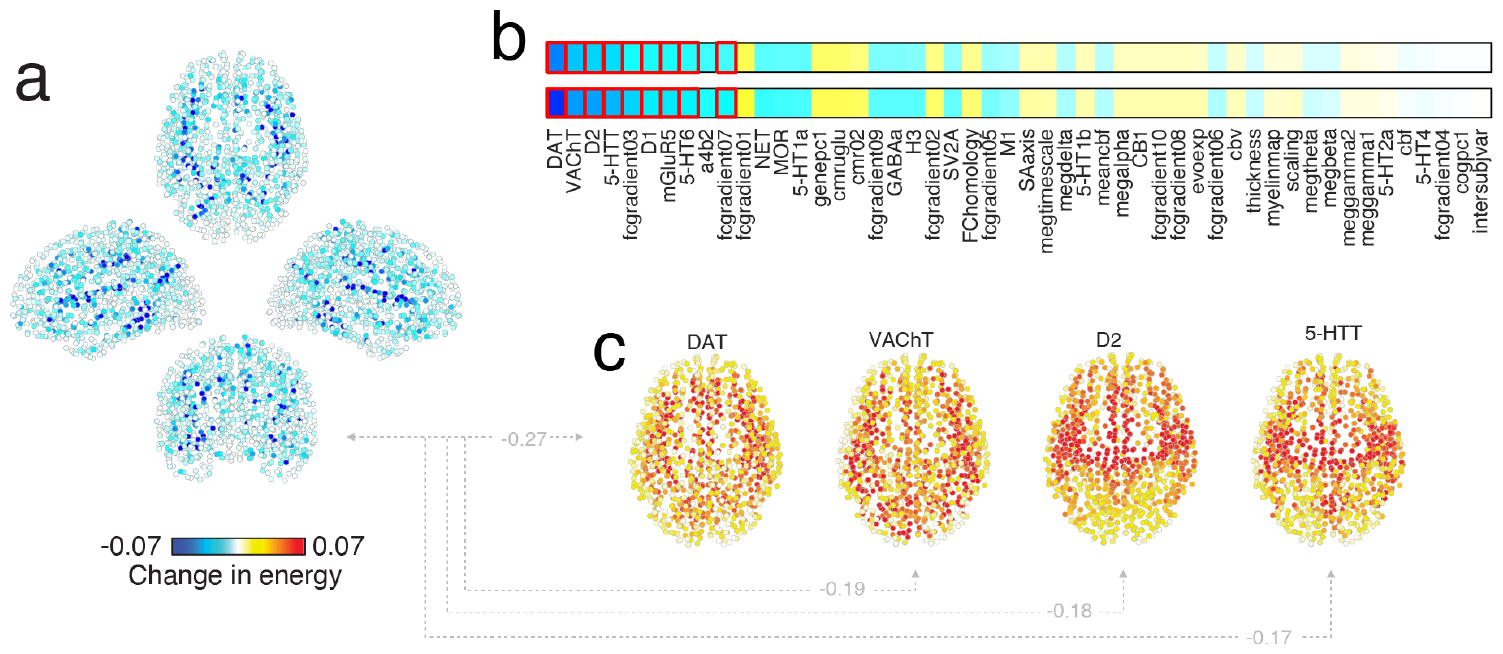
Regions whose energy increased under the spatial input strategy. In the main text we noted that a small subset of brain regions exhibited consistent increases in energy under the spatial input strategy compared to the local. (*a*) Spatial distribution of increases. (*b*) Correlation of the brain map from *a* with the annotations. Red outlines indicate statistically significant correlations. We show the annotation maps associated with the strongest statistically significant correlations in panel *c*.

**Figure S11.**
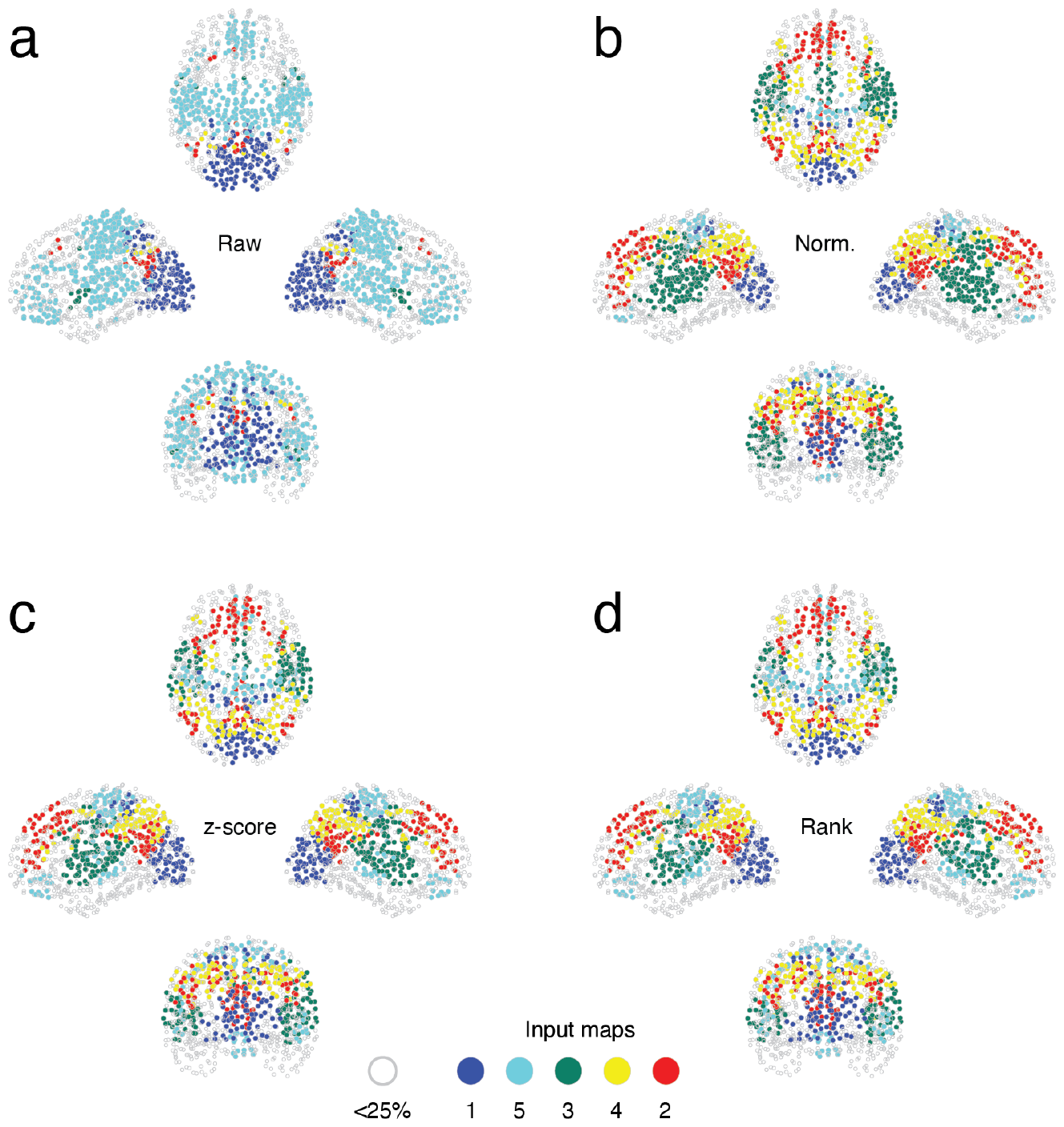
Dominance of input maps. In the main text we derived a series of brainwide input maps. Each map is associated with a specific input (control) signal; its elements correspond to how much of that signal should be delivered to each node. In the main text we focused on the top five maps, each of which appeared across many different control tasks. We also calculated for every node *i* its maximum input weight – that is, 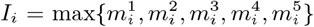 Here we show a series of “dominance maps”. That is, the index corresponding to the maximum input weight. We show four versions of the dominance maps, each of which we threshold so that the lowest quartile of maximum input weights are “greyed out”. (*a*) Dominance map using the raw (untransformed) input maps. (*b*) Dominance map after rescaling each map to the interval [0, 1] by subtracting the smallest input weight and dividing by the maximum value. (*c*) Dominance map after z-scoring the input maps. (*d*) Dominance map after rank-transforming each input map.

**Figure S12.**
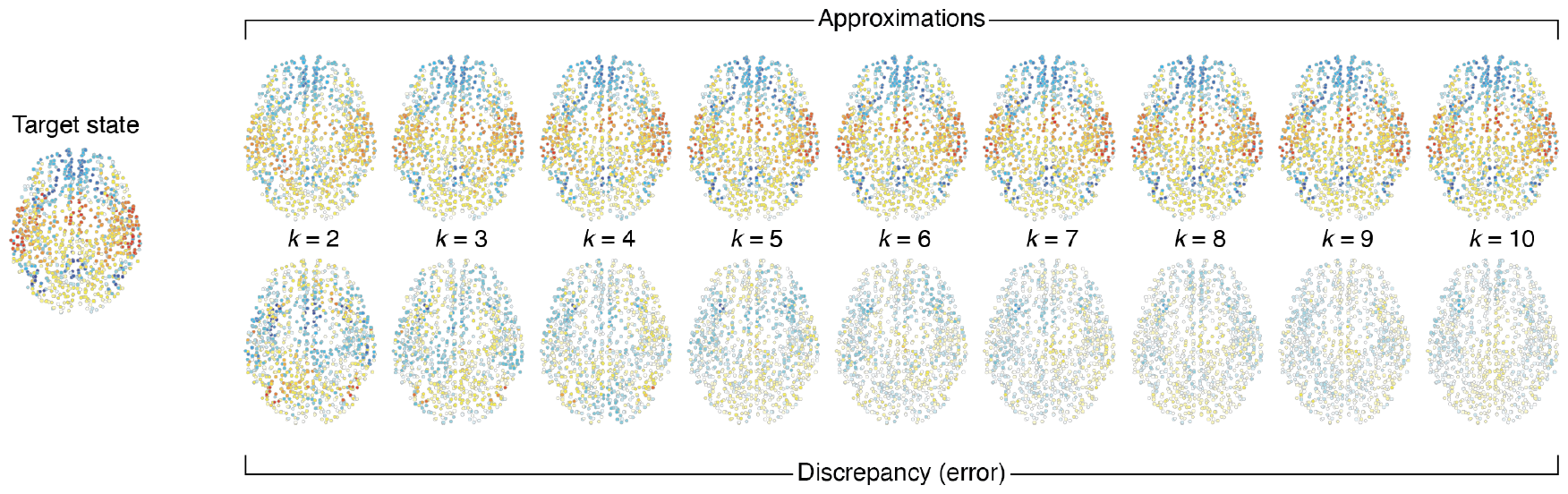
Example of approximation algorithm. On the left we show an example target state (state 1). The top row shows example approximations of the target state using *k* inputs instead of the *N* inputs usually required. The bottom row shows the error (approximation minus true state) for each *k* at a regional level.

